# Monitoring the directed evolution to a tripartite genome from a bipartite torradovirus genome

**DOI:** 10.1101/2024.03.20.585924

**Authors:** M. Turina, L. Nerva, M. Vallino, M. Ciuffo, B.W. Falk, I. Ferriol

## Abstract

We have previously shown that tomato apex necrosis virus that cannot express the RNA2-ORF1 protein (P21) is not able to systemically infect plant hosts but is not affected in cell autonomous aspects of virus replication/accumulation. Here we attempted to provide P21 *in trans* by co-agroinfiltrating the RNA2-ORF1 null constructs (a stop codon mutant and a deletion mutant) with a P21-expressing construct under control of the 35S promoter and containing the 5’ and 3’ UTRs of wild type (WT) RNA2. Such construct when co-agroinfiltrated with the stop codon mutant originates a WT bipartite virus through homologous recombination. More surprisingly, when co-agroinfiltrated with the P21 deletion mutant it cannot immediately complement the mutant, but it serendipitously originates a tripartite virus with an actively replicating P21-expressing RNA3 only after this replicating RNA3 accumulates deletions in a small region inside the original 3’-UTR provided by the cDNA clone. Such virus can be transmitted mechanically and by whiteflies, is competent for virion formation, and its RNA3 is encapsidated. The tripartite virus can be mechanically transferred for eleven generations without losing its infectivity or show major genomic rearrangements. Furthermore, mixing equal amounts of WT and tripartite virus inocula in the same leaf originated plants systemically infected only with the WT virus, showing that the tripartite virus has lower fitness than the WT. To our knowledge this is the first example of a stable virus evolving *in vitro* from bipartite to tripartite genomic structure from a synthetic construct in a plant virus.

## INTRODUCTION

Torradoviruses are newly emergent RNA plant viruses belonging to the family *Secoviridae* (Order *Picornavirales*) (Fuchs et al., 2022; Sanfaçon et al., 2009). The genus *Torradovirus* includes viruses of the species: *Torradovirus lycopersici* such as tomato torrado virus (ToTV) (Verbeek et al., 2007), *Torradovirus marchitezum* such as tomato marchitez virus (ToMarV) and tomato apex necrosis virus (ToANV) (Turina et al., 2007; Verbeek et al., 2008), *Torradovirus lactucae* including lettuce necrotic leaf curl virus (LNLCV) (Verbeek et al., 2014a), *Torradovirus carotae* including carrot torradovirus 1 (CaTV1) (Adams et al., 2014), *Torradovirus manihotis* including cassava torrado-like virus (CsTLV) (Carvajal-Yepes et al., 2014; Leiva et al., 2022), *Torradovirus cardiacae* motherwort yellow mottle virus (MYMoV) (Seo et al., 2015), *Torradovirus codonopsis* including codonopsis torradovirus A (Belete et al., 2021), and *Torradovirus cucurbitae* including squash chlorotic leaf spot virus (SCLSV) (Lecoq et al., 2016). Other tentative not yet assigned members are tomato chocolate virus (ToChV) (Verbeek et al., 2010), tomato chocolate spot virus (ToChSV) (Batuman et al., 2010), tomato necrotic dwarf virus (ToNDV) (Larsen et al., 1984; Wintermantel and Hladky, 2013), red clover torradovirus 1 (Koloniuk et al., 2018); physalis torrado virus (PhyTV) (Corrales-Cabra et al., 2021), fleabane yellow mosaic virus (FbYMV) (Alvarez-Quinto et al., 2022), burdock mosaic virus (BdMV) (Li et al., 2023), soybean torrado virus 1 (Rahman et al., 2023), potato rugose stunting virus (PotRSV) (Alvarez Quinto et al., 2023). Members of the genus *Torradovirus* are transmitted by whitefly (Verbeek et al., 2014b), aphid vectors (Rozado-Aguirre et al., 2016; Verbeek et al., 2017) and recent evidence indicates that psyllids can transmit PotRSV (Alvarez Quinto et al., 2023). Torradoviruses are single stranded positive-sense bipartite viruses that separately encapsidate each genomic RNA in isometric particles. The RNA1 (∼7 kb) has one open reading frame (ORF) and codes for a polyprotein which is proteolytically processed into specific proteins. The RNA2 (∼5 kb) has two open reading frames. The RNA2-ORF1 codes for a protein of approximately 21 kDa (P21) that is unique for torradoviruses and is required for systemic infection in tomato and *Nicotiana benthamiana* plants (Ferriol et al., 2018). The RNA2-ORF2 codes for a polyprotein which is processed by the 3C-like RNA1 protease into a movement protein and three capsid protein subunits (Ferriol et al., 2016a). The RNA1 is required for the replication of the RNA2 and for the RNA2 proteolytic processing of RNA2-encoded proteins (Ferriol et al., 2016a, 2016b).

Despite, the recent emergence of new torradoviruses, the molecular biology of torradoviruses has not been well characterized. Infectious cDNA clones are available for two tomato-infecting torradoviruses: ToTV (Wieczorek et al., 2015) and ToANV (Ferriol et al., 2016b). The role of torradovirus-encoded proteins in the infection cycle has been explored using different approaches such as expression vectors or infectious cDNA clones. Different genomic regions have been shown to be involved as determinants of pathogenicity such as the N-terminal region of the RNA1 of ToTV, the VP26 coat protein subunit and the 3A movement protein (Wieczorek et al., 2019; Wieczorek and Obrępalska-Stęplowska, 2016; Wieczorek Przemysław and Obrępalska-Stęplowska, 2016). Previously, we determined by reverse genetics the role of P21 in long distance movement in the infection cycle of ToANV (Ferriol et al., 2018). Recently, the RNA2-ORF1 protein of CsTLV was used to facilitate the mechanical inoculation of a potexvirus with low rate of mechanical inoculation in *Nicotiana benthamiana* plants (Jimenez et al., 2023) showing an interesting synergistic property. Although many functions of the protein have been characterized, we attempted to determine whether this protein is a cis- or a trans-active element. As part of further characterizing the functionality of the P21, in this work we designed an experimental system to test if providing P21 *in trans* was sufficient to complement ToANV RNA2-ORF1 functional mutants. Complementation of the ToANV RNA2-ORF1 protein *in trans* lead to unexpected results that yielded a functional and stable tripartite ToANV from the ToANV bipartite genome through specific deletions in the 3’ untranslated region (UTR). This new tripartite torradovirus was used to engineer a new ToANV-based GFP vector capable of infecting systemically *Nicotiana benthamiana* plants.

## RESULTS

### Providing P21 “*in trans*” to complement the p21 stop codon mutant results in delayed systemic infection requiring recombination events

Previously, we determined that the ToANV RNA2-ORF1 protein is required for systemic infection, but not for cell-to-cell movement, virion formation or replication in *N. benthamiana* plants (Ferriol et al., 2018). Such demonstration was based on two RNA2-ORF1 constructs that were not able to express the RNA2-ORF1 protein: R2-p21-sc and R2-Δp21 (Fig. 1 and Table 1). The R2-p21-sc construct contained three stop codons in the RNA2-ORF1 coding sequence, whereas R2-Δp21 had a deletion corresponding to a cDNA region coding for 190 amino acids (aas) (almost the entire cDNA corresponding to the RNA2-ORF1 protein). Agroinfiltration experiments in *N. benthamiana* plants using R2-p21-sc and R2-Δp21 constructs in combination with R1-WT construct resulted in replication in inoculated areas, but an inability to infect systemically *N. benthamiana* and tomato plants (Ferriol et al., 2018). In this work, we attempted to complement *in trans* the RNA2-ORF1 protein function by engineering a third construct that contained the RNA2-ORF1 protein coding sequence and the 5’- and 3’-UTRs of the RNA2-WT (R3-p21, Fig.1). Agroinfiltration assays of the R3-p21 construct in combination with R1-WT and R2-p21-sc constructs in *N. benthamiana* plants resulted in systemic infections, but there was a delay in symptoms appearance and virus presence with plants showing symptoms only 15 days post-agroinfiltration (dpa), contrary to what happens with ToANV-WT constructs (R1-WT+R2-WT) where most of *N. benthamiana* plants were systemically infected at 4 dpa (Table 1). This could imply a lack of direct and immediate complementation *in tran*s of the mutant. RT-PCR using specific primers to identify the R3-p21 and R2-p21-sc-derived RNAs (Table S1) performed on total RNAs obtained from upper non-inoculated leaves of *N. benthamiana* agroinfiltrated with R1-WT+R2-p21-sc+R3-p21, revealed that the third RNA provided *in trans* by R3-p21 through agroinfiltration was not present in the systemically infecting viral progeny. Furthermore, the nucleotide sequence of R2-p21-sc had reverted to the wild-type sequence, thus eliminating the stop codons in the viral progeny (data not shown). To estimate whether those changes in the viral progeny were due to recombination between the R2-p21-sc- and R2-p21-derived RNAs (and not reversions in the three stop codon position or contamination events), we generated the construct R3-p21-rc (Fig. 1): an R3-p21-derived construct with a silent mutation at the aa G^5^ (T to A at position 165; GGT to GGA accession KT756877) in the RNA2-ORF1 sequence that would allow us to monitor a possible recombination event. When we agroinfiltrated *N. benthamiana* plants with *A. tumefaciens* cells harboring R1-WT+R2-p21-sc+R3-p21-rc constructs, 5 out of 5 inoculated plants were systemically infected by 15 dpa. Sequencing of RT-PCR products corresponding to the mutated region of P21 from total RNAs of upper non-inoculated plants (three samples) confirmed the presence of the silent mutation in the sequence of the RNA2 demonstrating that a recombination event between the RNA2 derived from R2-p21-sc containing the stop codon mutations and the RNA expressed from R3-p21-rc gave rise to the restored progeny (Supplementary Fig. 1).

**Figure 1.**
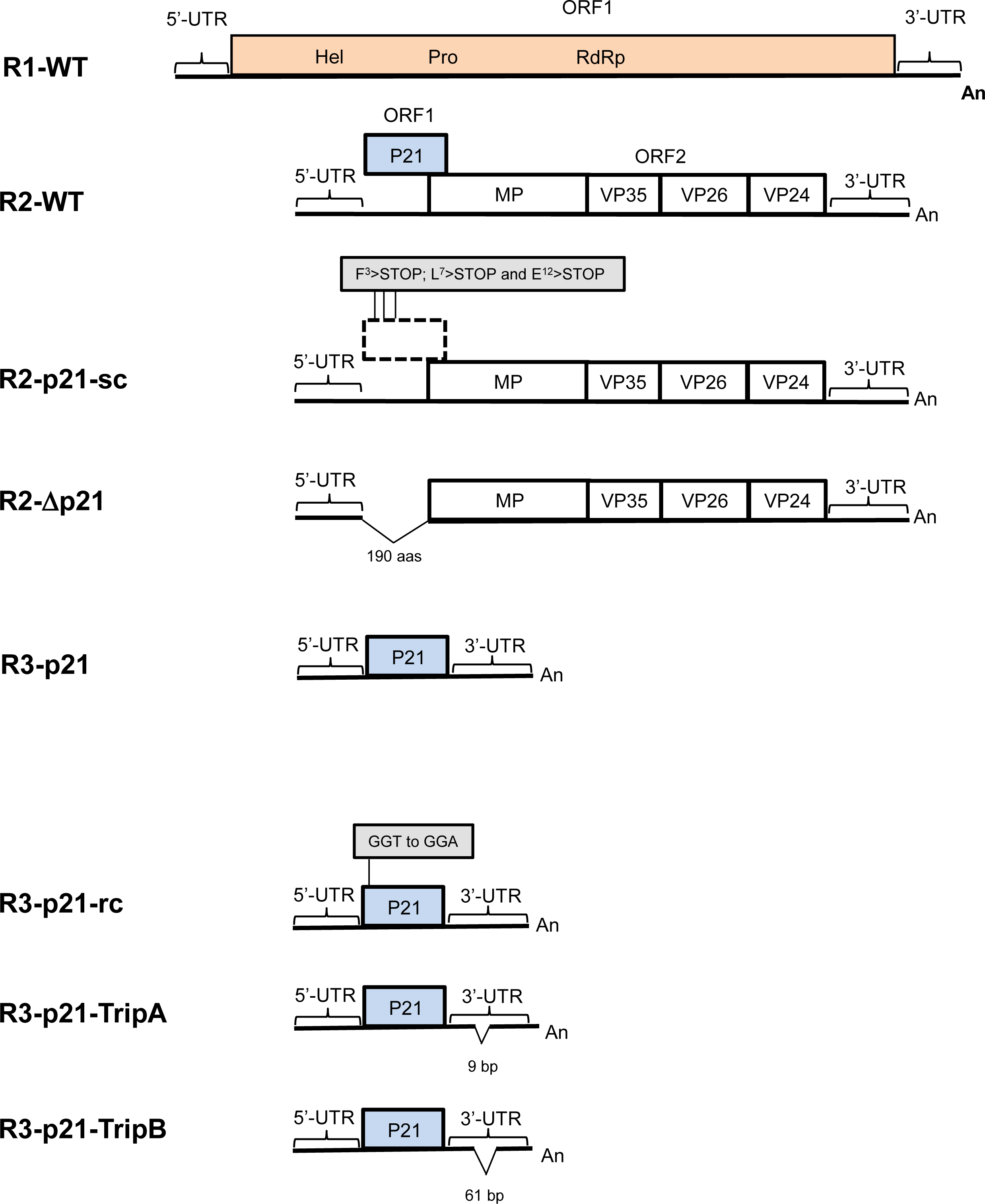
Constructs used in this study. Diagram showing the constructs used in this study. The construct pJL89-M-R1 (R1-WT), pJL89-M-R2 (R2-WT), R3-p21(Ferriol et al., 2016b) and the series of R3-p21 3’-UTR deletion mutants are shown. Horizontal lines represent the RNA genome and boxes indicate open reading frames (ORFs). The RNA2-encoded proteins are: RNA2-ORF1 protein (P21, represented with a blue box), movement protein (MP) and the three capsid proteins (VP35, VP26, VP24). The 5’ and 3’ untranslated regions (UTRs) are shown as black lines highlighted by a horizontal square bracket. The polyadenylated tail is shown at the 3’ end of the RNA (An). The positions of the stop codons introduced in R3-p21-sc are highlighted in a grey box. Details of the ToANV RNA2-derived deletion mutants are indicated.

**Table 1.** Trans complementation experiments to assess the ability of a putative RNA2-ORF1 expressing construct to supplement the activity *in trans* of the RNA2-ORF1 knockout mutants designed in tomato apex necrosis RNA2.

| Inoculated constructs<sup>a</sup> | Experiment | 4dpa<br>LOCAL<sup>b</sup> | 4dpa<br>SYSTEMIC<sup>b</sup> | 7 or 8dpa<br>SYSTEMIC<sup>b</sup> | 15 or 16dpa<br>SYSTEMIC<sup>b</sup> |
| --- | --- | --- | --- | --- | --- |
| R1-WT+R2-WT | Exp1 | 2/2 | 2/2 | 2/2 | N/A |
|  | Exp2 | 2/2 | 5/5 | N/A | N/A |
|  | Exp3 | N/A | 13/15 | 15/15 | 15/15 |
| R1-WT+R2-p21-sc | Exp1 | 2/2 | 0/2 | 0/2 | 0/2 |
|  | Exp2 | 5/5 | 0/5 | 0/5 | 0/5 |
|  | Exp3 | N/A | N/A | 0/15 | 0/15 |
| R1-WT+R2-p21-sc+<br>R3-p21 | Exp1 | 2/2 | 0/2 | 0/2 | 2/2 |
|  | Exp2 | 5/5 | 0/5 | 0/5 | 5/5 |
|  | Exp3 | N/A | N/A | 0/15 | 10/15 |
| R1-WT+R2-Δp21 | Exp1 | 2/2 | 0/2 | 0/2 | 0/2 |
|  | Exp2 | 5/5 | 0/5 | 0/5 | 0/5 |
|  | Exp3 | N/A | N/A | 0/15 | 0/15 |
| R1-WT+ R2-Δp21+<br>R3-p21 | Exp1 | 2/2 | 0/2 | 0/2 | 1/2 |
|  | Exp2 | 5/5 | 0/5 | 0/5 | 1/5 |
|  | Exp3 | N/A | N/A | 0/15 | 2/15 |

<sup>a</sup> Constructs are those described in Figure 1 and in the text.
<sup>b</sup> The numbers in the these columns represent the numbers of infected plants/number of tested plants present in the experiment; DPA= days post agroinfiltration. Tests to confirm virus infection were carried out through western blot (for Exp1 and Exp2) and by DAS-ELISA for Exp3. N/A= test not carried out for the specific experiment. Local and Systemic, refers to inoculated leaves and upper un-inoculated leaves respectively

### Providing P21 “*in trans*” to complement the R2-Δp21 deletion mutant results in occasional delayed systemic infection with a tripartite virus progeny

We next attempted to complement the RNA2-ORF1 deletion mutant (R2-Δp21) construct with our R3-p21 construct. *N. benthamiana* plants agroinfiltrated with R1-WT+R2-Δp21+R3-p21 only became occasionally infected after 15 dpa (Table 1). Northern blot analysis of total RNAs from upper non-inoculated leaves of plants agroinfiltrated with the R2-Δp21 and R3-p21 constructs showed the presence of three RNAs of positive and negative sense, with sizes putatively corresponding to the RNA1-, RNA2-Δp21- and RNA3-p21-derived replicating RNAs (Fig. 2). Presence of relatively abundant –strand RNA corresponding to RNA3 demonstrates that complementation is not purely through systemic movement of the third RNA provided locally through agroinfiltration, but through active replication of this third RNA. Sequencing RT-PCR products for RNAs 1, 2 and 3 from total RNAs from the upper, non-inoculated leaves showed the absence of nucleotide changes in the R2-Δp21-derived RNAs, where the P21 coding sequence was deleted. However, there were deletions in the 3’-UTR of R3-p21-derived third RNA. Deletions of 6-, 9- and 61-nt were present in the 3’-UTR of R3-p21-derived RNAs; these deletions were obtained from three different *N. benthamiana* plants from three different experiments (Fig. 3).

**Figure 2.**
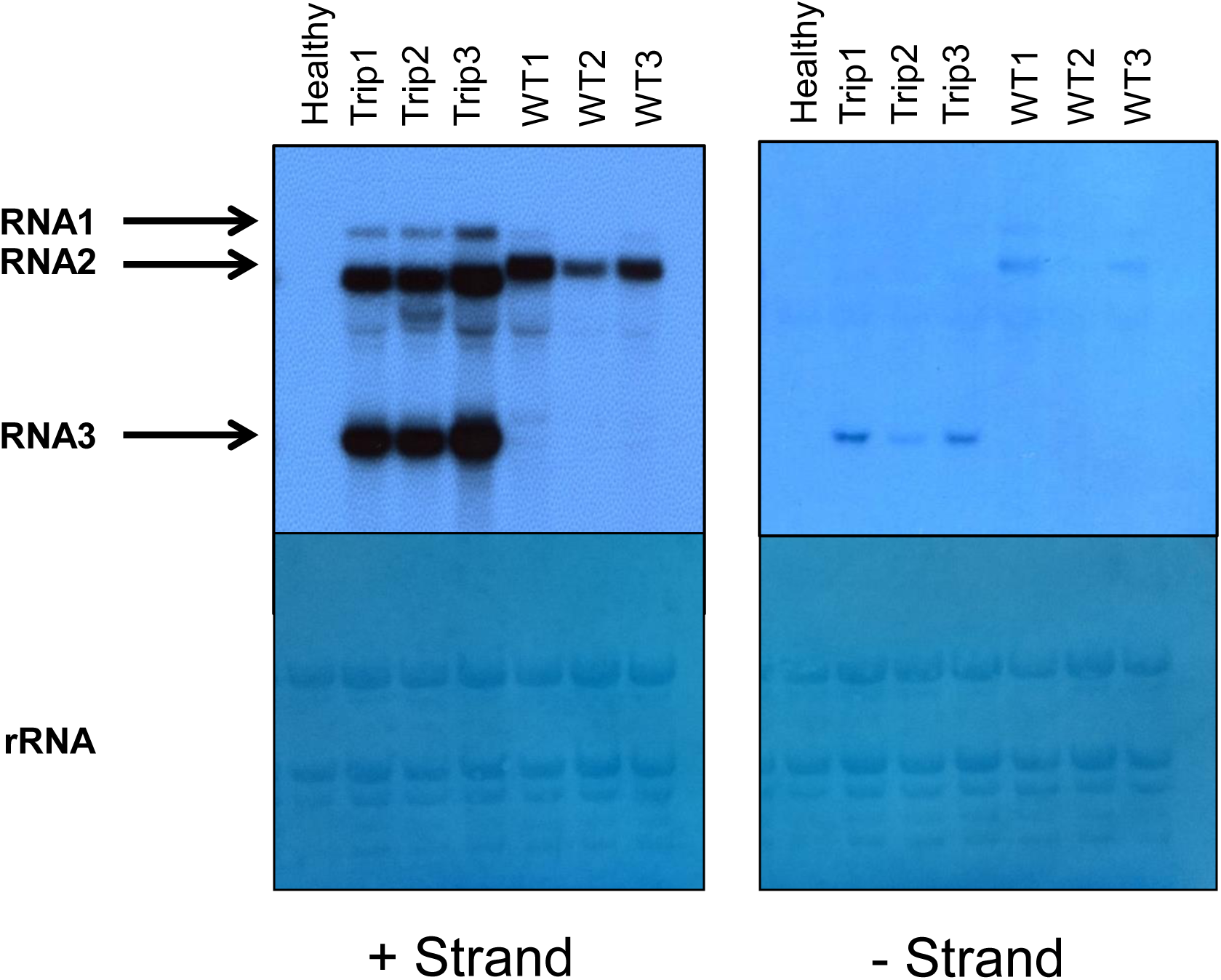
Attempts at complementation of a p21 deletion mutant providing p21 “*in trans*” results in systemic infection with a replicating tripartite virus. Northern blot analysis using plus strand (left panels) and minus strand (right panels) hybridizing probes, designed in the 3’-UTR region common to RNA1 and RNA2 of tomato apex necrosis virus reveal that in extracts from systemically infected plants that were inoculated with R1-WT+R2-Δp21+R3-p21 (called Trip1, Trip2 and Trip3) we can detect both plus strand and minus strand RNAs corresponding to a smaller genomic RNA of the size predicted based on R3-p21 transcript size. WT1, WT2, and WT3 are lanes loaded with total RNA from plants inoculated with R1-WT+R2-WT constructs. Ribosomal RNA loading (rRNA) can be compared in the two bottom panels representing the same membranes of the top panels stained with methylene blue. Healthy, correspond to total RNA extracted from mock inoculated plants. Arrows point to RNA1, RNA2 and RNA3. Note that RNA2 size in the Trip1-3 samples carries the deletion provided in the infectious clone (smaller size).

**Figure 3.**
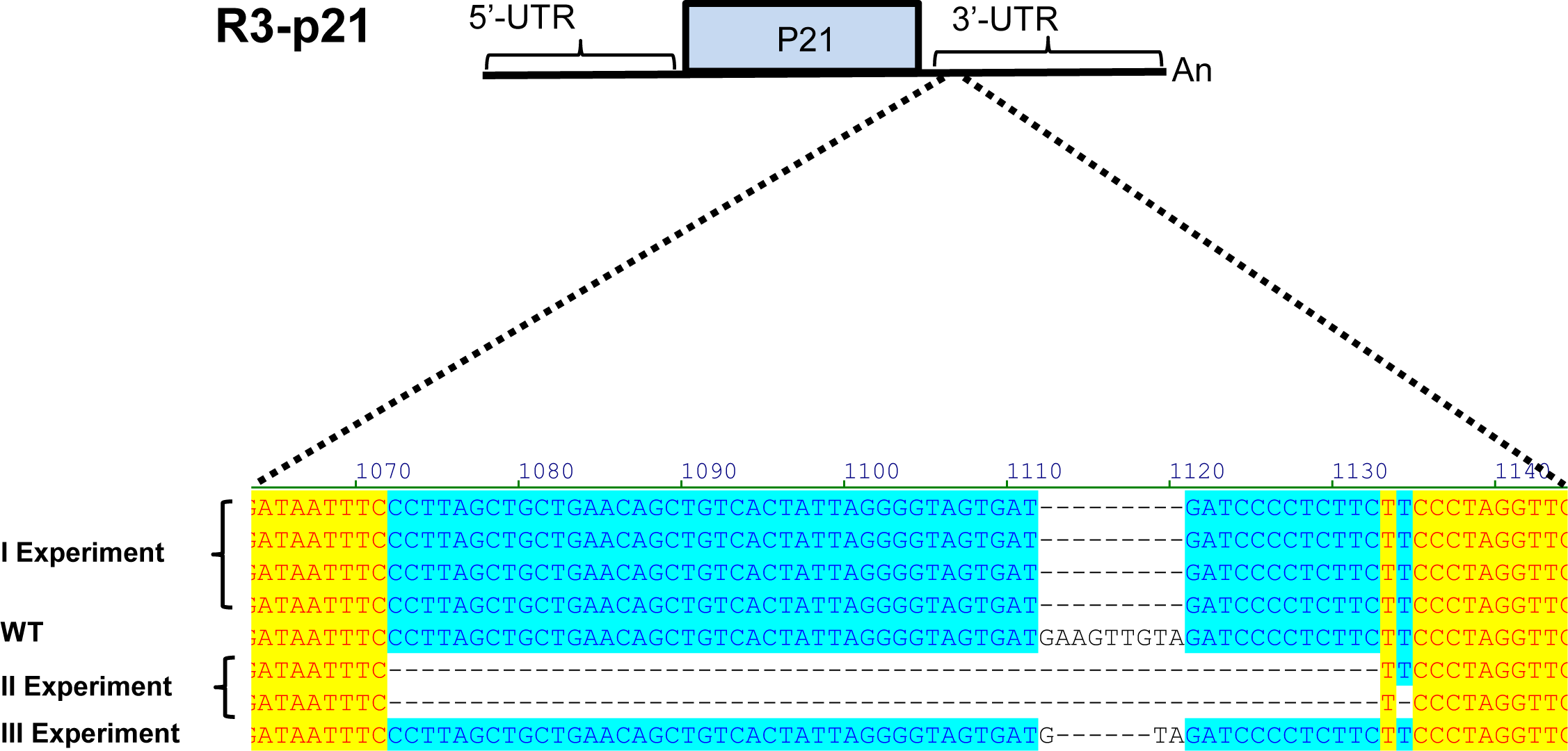
The third replicating RNAs in tripartite tomato apex necrosis viruses carry specific deletions in the 3’-UTR. For each of the three complementation experiments described in Table 1 that yielded a tripartite systemic virus infection (R1-WT+R2-Δp21+R3-p21), we performed a full-length Sanger sequence of the putative RNA3, for one systemically infected plant in each of the three experiments. Sequence alignments reveal that all three tripartite viruses carry a specific deletion in the same region of the 3’UTR, each characterized by a different size ranging from 6 to 61. Numbers in the alignment refer to the sequence of the cDNA clone provided *in trans*, schematically reproduced in the upper part of the figure. Parenthesis include different cDNA clones sequenced from the same systemically infected plant.

To investigate whether these deletions have any effect on the secondary structure of the 3’-UTR of each of the derived RNAs, we compared the predicted secondary structures using the MFold program. The predicted RNA secondary structures of the 3’-UTRs of R3-p21, R2-WT and R3-p21-6bp-deletion showed similar RNA secondary structures. However, R3-p21-9bp-deletion and R3-p21-61bp-deletion derived RNAs had different secondary structures when compared with R3-p21 and R2-WT (Supplementary Fig. 2). In addition, multiple nucleotide alignment of the RNA1 and RNA2 3’-UTRs of different ToMarV isolates showed that these deletions are not present in other ToMarV naturally occurring isolates (Supplementary Fig. 3).

### Biological characterization of the newly derived tripartite torradovirus

We engineered two R3-p21-derived constructs with the deletion in the 3’-UTR region which we obtained in experiment I and experiment II (9-bp and 61-bp, respectively) and the resulting constructs were named R3-p21-TripA and R3-p21-TripB (Fig. 1). To determine whether these constructs were indeed able to immediately replicate as such without further mutations and mimic the systemic infection of the WT virus, we agroinfiltrated four *N. benthamiana* plants with the two ToANV “tripartite viruses”, hereafter referred as ToANV-TripA (R1-WT+ R2-Δp21+R3-p21-TripA) and ToANV-TripB (R1-WT+ R2-Δp21+R3-p21-TripB). As a positive control we also agroinfiltrated 5 *N. benthamiana* plants with ToANV WT constructs (R1-WT+R2-WT). At 4 dpa, all plants showed typical symptoms of infection and we confirmed the presence of the three RNAs in the derived-progeny of ToANV-TripA and ToANV-TripB by northern blot analyses (Fig. 4) showing that indeed only after the replication-dependent 3’-UTR region deletions, the P21 expressing construct can complement *in trans* the loss of function of RNA2 P21 knock out. Our results showed that both tripartite viruses had three replicating RNAs corresponding to their expected sizes.

**Figure 4.**
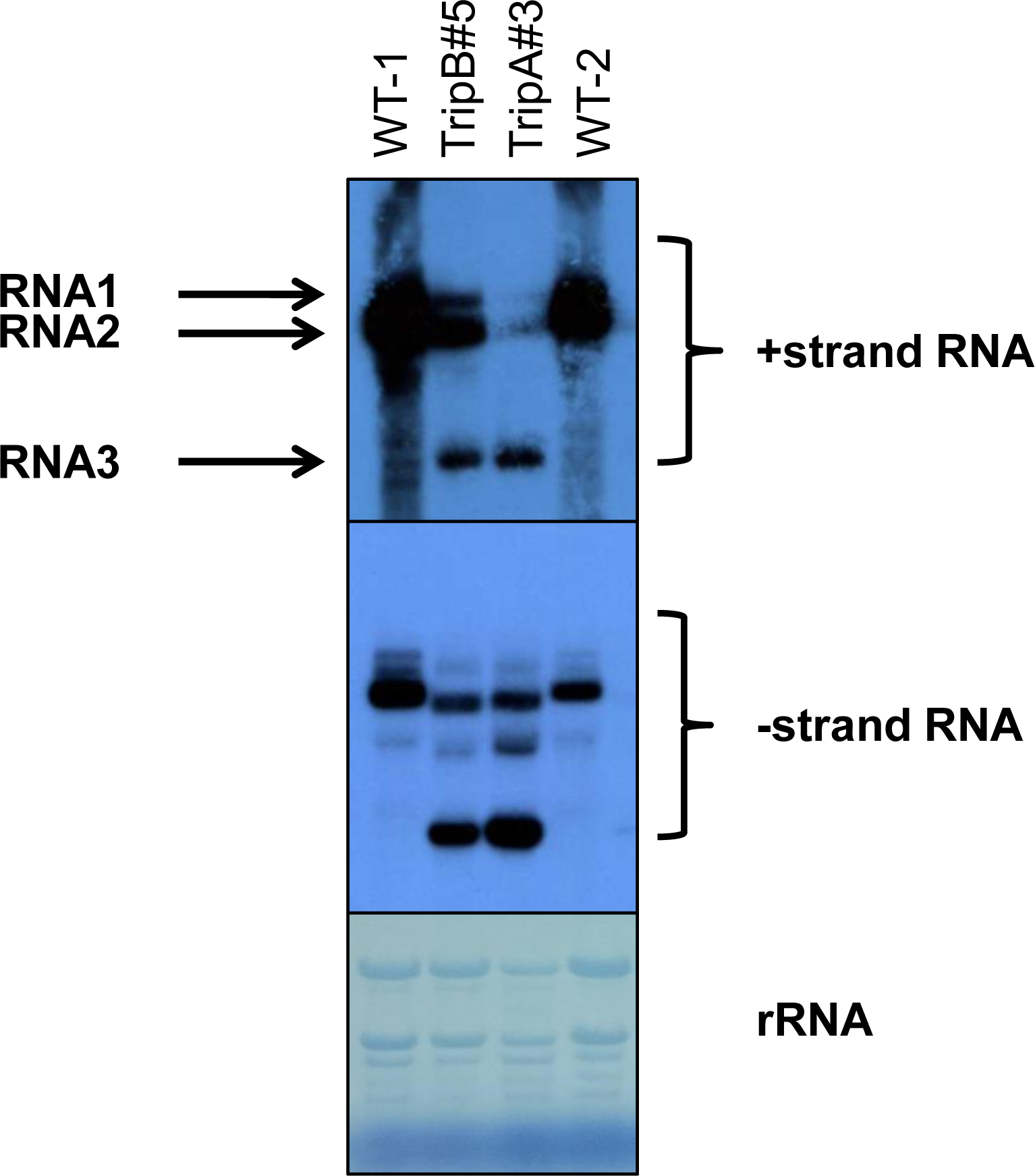
Agroinoculation of the artificially created tomato apex necrosis tripartite virus results in efficient systemic infection. Northern blot analysis using plus strand (upper panel) and minus strand (lower panel) hybridizing probes, designed for the 3’-UTR regions common to RNA1 and RNA2 of tomato apex necrosis virus reveal that actively replicating tripartite virus can be detected in systemically infected leaves harvested four days post agroinfiltration. WT-1 and WT-2 are lanes loaded with total RNA extracts from leaves from two distinct systemically infected plants inoculated with R1-WT+R2-WT 4 days before collecting the samples. TripA#3 and TripB#5 are extracts of systemically infected plants harvested four days post agroinfiltration with the following constructs: R1-WT+R2-Δp21+R3-p21-TripA and R1-WT+R2-Δp21+R3-p21-TripB, respectively. Ribosomal RNA loading (rRNA) can be compared in the bottom panel representing the same membranes of the top panels stained with methylene blue. Arrows point to RNA1, RNA2 and RNA3. Note that RNA2 size in the TripA#3 and TripB#5 samples carry the deletion provided in the infectious clone (smaller size).

Defective mutants arise many times from the populations of multipartite viruses, but the biological properties and fitness of these variants is often different from the wild-type virus (García-Arriaza et al., 2004). Thus, we decided to focus on one of our tripartite viruses (ToANV-TripB) and determine if its biological properties were comparable to ToANV WT and test Koch’s postulates for the ToANV-TripB. First, we compared the host range of the bipartite (ToANV WT) and tripartite (ToANV-TripB) viruses, agroinfiltrating different hosts (Table 2). Our results showed that ToANV WT and the tripartite virus ToANV-TripB had the same biological host range for the hosts tested providing the infection through agroinfiltration. Symptoms were similar in the plant species tested except for tomato, where systemic symptoms were milder for the tripartite virus (Supplementary Fig. 4). Second, we checked whether the ToANV-TripB was able to be mechanically inoculated during several passages and whether the ToANV-TripB was maintained over such passages. We performed eleven passages in *N. benthamiana* plants and evaluated the viral progeny after these passages by northern blot analyses (Fig. 5). Our results showed that after eleven passages the tripartite virus ToANV-TripB was maintained in the population without major genomic rearrangements; the sequence of these viruses passaged mechanically was determined by next generation sequencing (NGS) and is discussed below. When mechanical inoculation of ToANV-TripB was attempted from tomato plant extracts to tomato plants (susceptible cv York), we were not able to reproduce infection, showing in this case a different outcome compared to ToANV WT virus.

**Figure 5.**
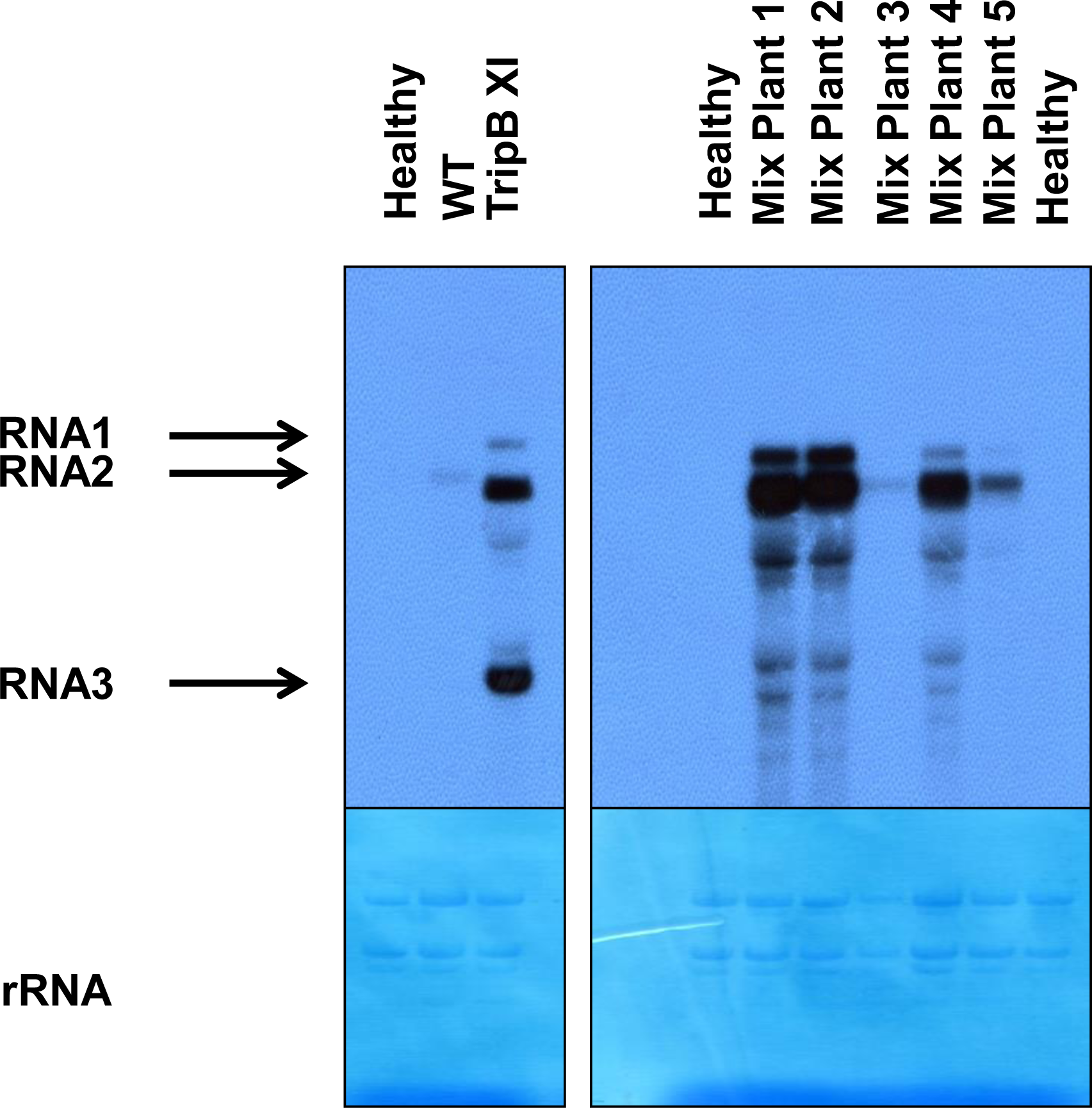
Tomato apex necrosis tripartite virus can be maintained through mechanical inoculation for eleven passages but is quickly overcome by the wild type virus in co-inoculation experiments. The left panels display samples of *Nicotiana benthamiana* plants mechanically inoculated serially with virus originated from WT infection (R1-WT+R2-WT) and from TripB infection (R1-WT+R2-Δp21+R3-p21-TripB). The WT sample was collected at time of infection when recovery of symptoms occurred (20 dpi), while the TripB XI represents a sample collected at the eleventh mechanical inoculation passage in a symptomatic stage (5 dpi). The right-side panels display northern blot analysis of leaf extracts from systemically infected plants harvested 7 days post agroinfiltration with a mix of equal amounts of *A. tumefaciens* harboring the R2-WT clone and the R2-Δp21+R3-p21-TripB clones. Each lane is loaded with a different plant from the same experiment (Mix Plant1 to Mix Plant 5). Northern blot was carried out using plus strand hybridizing probes, designed in the 3’-UTR region common to RNA1 and RNA2 of tomato apex necrosis virus. Ribosomal RNA loading (rRNA) can be compared in the bottom panels representing the same membranes of the top panels stained with methylene blue. Arrows point to RNA1, RNA2 and RNA3. Healthy represents extracts from mock inoculated plants.

**Table 2.** Host range of plants agroinfiltrated with ToANV WT or ToANV Trip B 14 days post agroinfiltration (dpa).

| Host | Constructs<sup>a</sup> |  |
| --- | --- | --- |
|  | ToANV WT <sup>b</sup> | ToANV Trip B <sup>b</sup> |
| *Nicotiana clevelandii* | 2/2 | 2/2 |
| *N. rustica* | 0/2 | 0/2 |
| *N. tabacum* White Burley | 2/2 | 2/2 |
| *Solanum lycopersicum* cultivar York | 2/2 | 2/2 |
| *Capsicum annuum* cultivar Quadrato D'asti | 0/2 | 0/2 |
| *N. megalosiphon* | 2/2 | 2/2 |
| *Datura stramonium* | 2/2 | 1/2 |
| *N. benthamiana* | 2/2 | 2/2 |
| *N. glutinosa* | 2/2 | 2/2 |
| *N. occidentalis* | 2/2 | 1/2 |

<sup>a</sup> Constructs agroinfiltrated in different host plants: ToANV WT (R1-WT+R2-WT) or ToANV TripB mutant (R1+R2 R2Δp21+R3-p21-TripB).
<sup>b</sup> Number of plants infected divided by the number of total plants agroinfiltrated after 14 dpa. ToANV infection was detected by double antibody sandwich-enzyme-linked immunosorbent assay (DAS-ELISA) carried out on extracts from the upper un-inoculated leaves of different hosts.

We then performed a competition assay by agroinfiltration of both viruses (ToANV-WT and ToANV-TripB) in equal amounts based on the OD values of the *A. tumefaciens* cells used for infiltration in *N. benthamiana* plants. Systemically infected plants were analyzed by northern blot 7 dpa in order to verify the possible presence of the tripartite virus together with the wild type. The amount of viral RNA that we detected in each systemically infected plant differed among the various replicates, but no evidence of the tripartite virus alone or in mixed infection was obtained (Fig. 5). Therefore, in a mixed infection the WT virus quickly outcompetes ToANV-TripB. Finally, we checked the whole genomic sequences of ToANV-WT and ToANV-Trip B virus infections looking at nucleotide variants of reads mapping on the two distinct genomes with their associated reference sequences (those of the infectious clones) (Supplementary Table 2 to 4). A number of nucleotide polymorphisms are present in specific positions, some of them appearing in all the different samples, but there is not a single specific change in RNA1 and RNA2 that is associated to the presence of a third replicating RNA.

### The tripartite virus yields virions that encapsidate RNA3

We then proceeded to investigate if the properties of ToANV-TripB virus were similar to ToANV WT in relationship to virion formation and RNA encapsidation. From systemically infected *N. benthamiana* plants we harvested equal amounts of leaves and purified virions according to a protocol previously described in detail (Turina et al., 2007). After sucrose gradient centrifugation, pure virion preparations of both viruses were obtained: SDS-PAGE analysis shows the three capsid proteins present in comparable amounts between ToANV WT and ToANV-TripB purified preparations (Fig. 6A) showing that ToANV-TripB also originates stable virions. We then proceeded to investigate if RNA3 was also encapsidated, and not simply moved systemically as free RNA. Northern blot analysis of post sucrose gradient purified virion preparations showed the presence of RNA3 (Fig. 6B). Transmission electron microscopy (TEM) observation of the preparations showed virions of similar size and shapes with a higher percentage of stain-penetrated empty (or partially full) particles (Fig. 6C).

**Figure 6.**
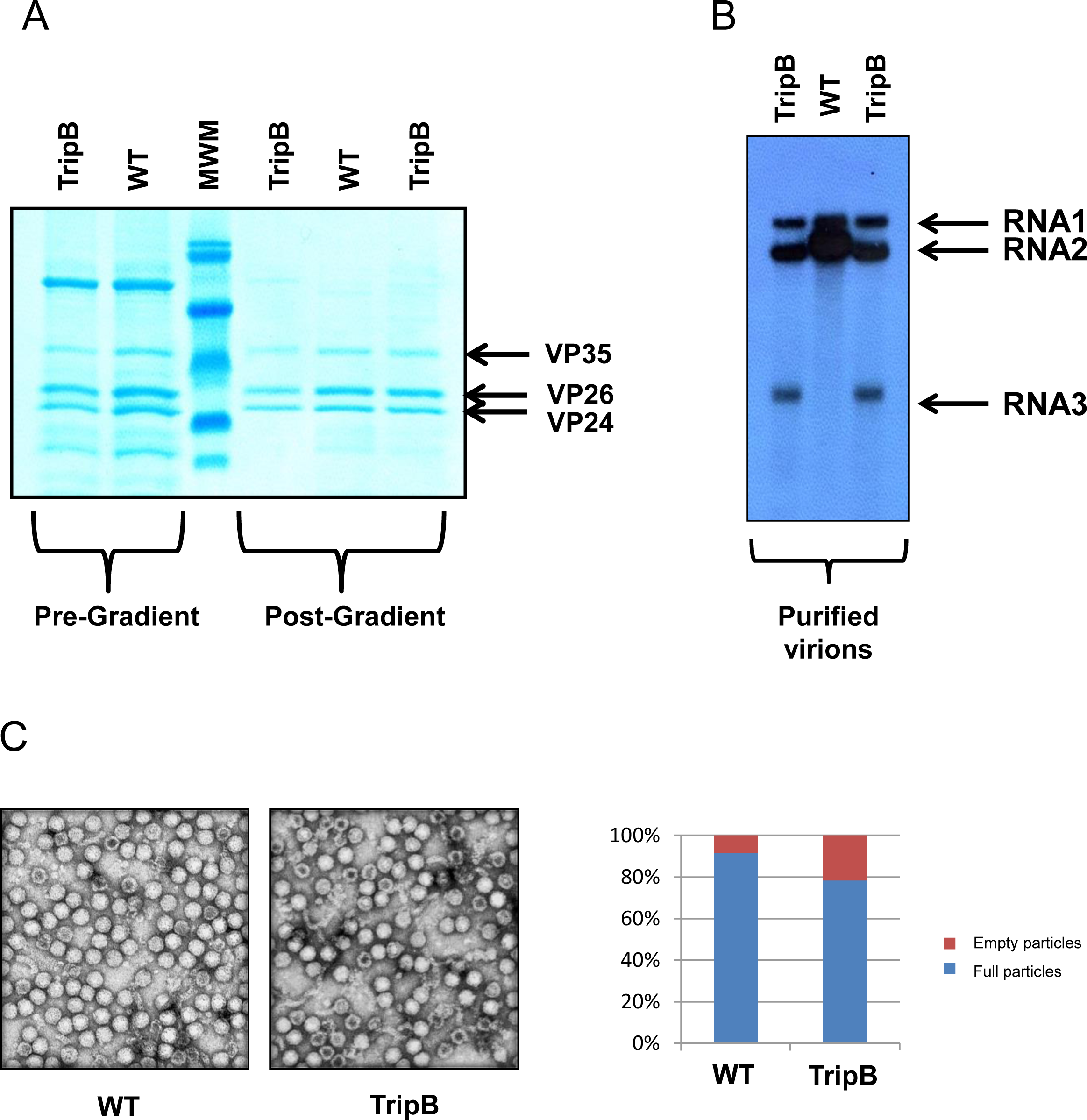
Tomato apex necrosis tripartite virus forms stable virions and encapsidates the third genomic RNA. Panel A displays Coomassie stained SDS-PAGE of samples from purified WT virions and TripB virus pre- and post-gradient. The arrows point at the three proteins that are present in purified virions (VP35, VP26 and VP24). MWM = molecular weight marker. Panel B displays northern blot analysis of RNA extracted from equal amount of the same virus purification displayed in panel A. Arrows in this panel point to the position of the bands representing the three genomic RNAs. Panel C displays representative pictures of virus purifications taken with the transmission electron microscope (TEM). The right-end graph shows the ratio of full and empty particles in each of the two virus purifications.

### The tripartite virus can be transmitted in controlled environment by whiteflies

Finally, to verify whether the ToANV-Trip B tripartite virus is competent for whitefly transmission, we performed *Trialeurodes vaporariorum* transmission experiments of the tripartite virus using the same acquisition times and inoculation access times as in previous experiments (Ferriol et al., 2016b). We used as acquisition host *N. benthamiana* systemically infected leaves 7 days post-agroinfiltration, whereas for inoculation we used both *N. occidentalis* and *Physalis floridana* plants. Using whiteflies left to acquire on WT infected leaves we could obtain successful transmission in 4 out of 6 inoculated *N. occidentalis* plants and 2 out of 5 *P. floridana* plants. When whiteflies were left to acquire on ToANV-TripB-infected plants 2 plants out of 6 *N. occidentalis* resulted systemically infected, whereas none of the three *P. floridana* plants were infected. We extracted RNA from a subset of plants showing systemic symptoms and a northern blot was carried out to reveal the presence and sizes of the viral RNAs (Supplementary Fig. 5): as expected the plant originated from whitefly which had acquired from ToANV WT RNA, showed a full-length RNA2, whereas the two samples extracted from plants inoculated with whiteflies which had acquired on ToANV-TripB-infected plants showed the presence of the RNA2 deleted form, and the RNA3. Overall, these results demonstrated that ToANV-TripB can be transmitted by its whitefly vector *T. vaporarorium*.

### The tripartite virus can be used to generate a new GFP expressing vector able to systemically infect *N. benthamiana* plants

Previously we have assembled a ToANV derived vector to express GFP engineered with predicted proteolytic sites between the MP and VP35 proteins on the RNA2 (Ferriol et al., 2016b). Such vector could not be used to monitor systemic infection since GFP could only be detected in inoculated tissues and virus systemic movement was impaired. Here, based on the occurrence of the tripartite ToANV-TripB we envisioned a new strategy to express GFP: our RNA3 expressing clone contains a second AUG just before the carboxyterminal of P21 which is the start codon of RNA2-ORF2. We therefore engineered a restriction site to insert in frame to this second AUG the GFP coding sequence originating the construct R3-p21-TripB-GFP (Fig. 7A) therefore making the RNA3 bicistronic. Upon agroinfiltration of *N. benthamiana* plants with R1-WT, R2-Δp21 and R3-p21-TripB-GFP we observed abundant GFP accumulation in the agroinfiltrated area 3 dpa, and by 6 dpa some enlarged foci in the un-inoculated leaves were also distinguishable. Also, some GFP-expressing infected leaf primordia in the germinating apex were easily distinguishable (Fig. 7B). No GFP expression was observed in the leaves corresponding to ToANV WT infection. This new tool will be pivotal in trying to fine-tune the role of specific P21 domains in long-distance spread elucidated through alanine scanning mutagenesis.

**Figure 7.**
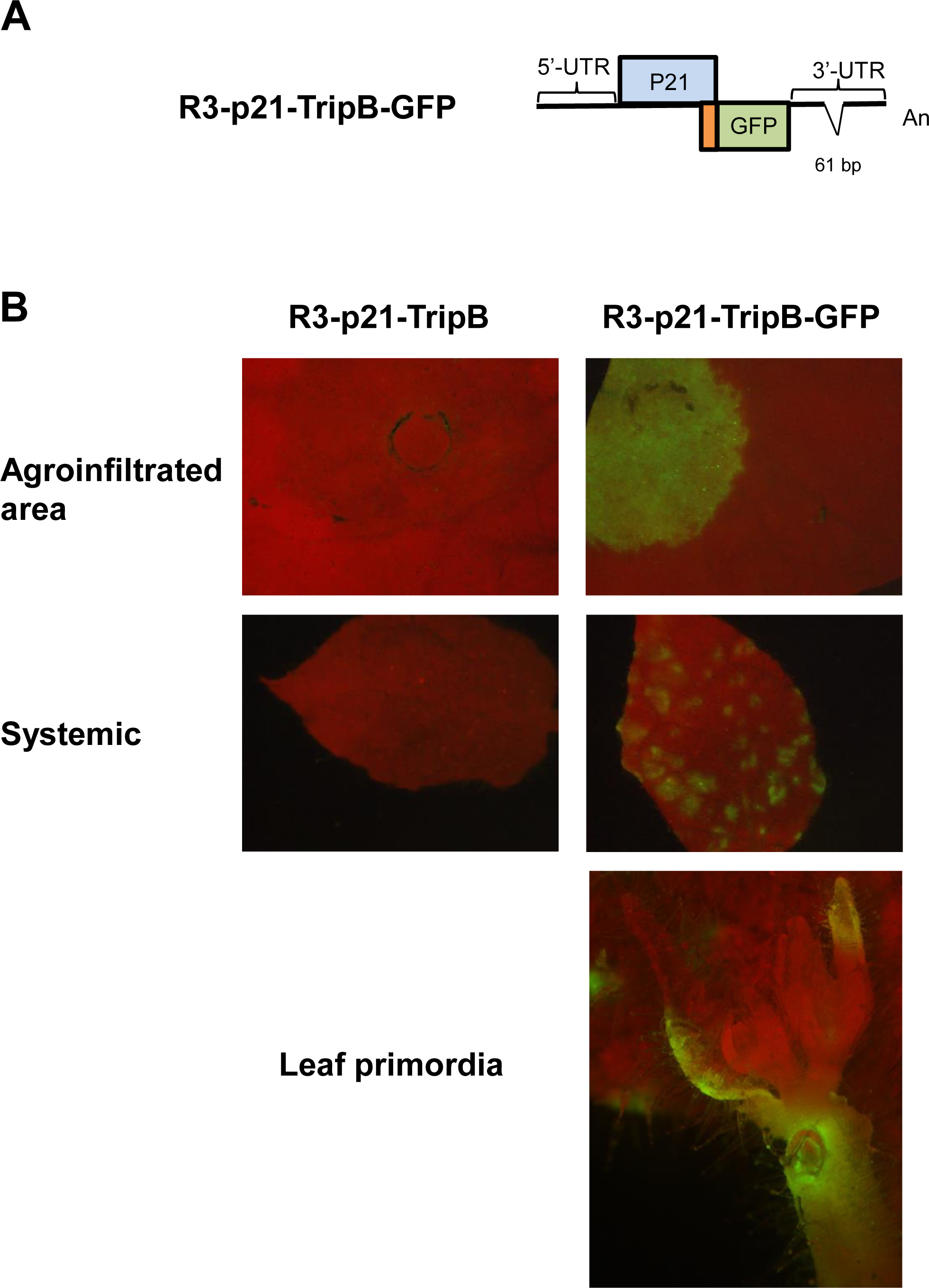
The tripartite construct allows to derive a new GFP viral vector able to infect *Nicotiana benthamiana* systemically. Panel A depicts the R3-p21-TripB-GFP construct showing the fusion between the initial amino acids of RNA2-ORF2 fused to GFP. Panel B displays *N. benthamiana* leaves observed under a fluorescence stereo microscope, with fluorescent light used to monitor GFP expression. Green fluorescence corresponding to GFP expression is observed both locally and systemically when the new GFP vector (R1-WT+R2-Δp21+R3-p21-TripB-GFP, labelled as R3-p21-TripB-GFP) is agroinfiltrated in *N. benthamiana* plants. Only red chlorophyll autofluorescence can be visualized in plants agroinfiltrated with R1-WT+R2-Δp21+R3-p21-TripB (labelled as R3-p21-TripB), used as negative control.

## DISCUSSION

One of the most intriguing peculiarities of members of the genus *Torradovirus* is the presence of an ORF in RNA2 upstream of the polyprotein (RNA2-ORF1, which codes for the protein P21), which is not present in other members of the family *Secoviridae*. Previously, we demonstrated that P21 protein is required for systemic infection in *N. benthamiana* and tomato plants, but not for cell-to-cell movement, virion formation or replication (Ferriol et al., 2018). To better characterize the role of the RNA2-ORF1 protein in the infection cycle of ToANV in plants, we wanted to test if P21 is functional *in trans* on RNA2 that cannot express it: for this purpose we provided the P21 protein “*in trans*” to two different RNA2-ORF1 mutants: one with stop codons inserted downstream of the AUG and one with an almost complete deletion of the coding sequence, R2-p21-sc and R2-Δp21, respectively. The complementation construct (R3-p21) included the intact 5’-UTR and 3’-UTR of RNA2 to possibly allow a replicative form of the construct, and not simply rely on 35S-dependent expression for P21 accumulation, which would occur without the original viral 5’- and 3’-UTRs. We have direct evidence that this construct expresses P21 protein *in vivo*, since we have previously shown by western blot analysis that the exact same construct with an engineered FLAG peptide at the carboxy terminal accumulates the fusion protein in agroinfiltrated leaves (Ferriol et al., 2018). Agroinfiltration of R2-p21-sc in combination with R3-p21 construct resulted in a delayed yet efficient systemic infection in *N. benthamiana* plants but all the viral progeny no longer had the inserted stop codons, but had a WT sequence RNA2. We demonstrated that this change is originated from recombination with R3-p21 derived RNA that has extensive region of homology in the coding region between the two constructs. To further delineate if the RNA2-ORF1 protein could provide its function “*in trans”*, we provided the R3-p21 construct in combination with R2-Δp21 construct and the outcome was the occasional (but reproducible in three experiments out of three performed) delayed infection with a tripartite virus with three different types of deletions in the RNA3 3’-UTR each obtained from different experiments. Overall, our approach showed that simple expression/replication *in trans* of a third RNA expressing P21 with intact 3’-UTR does not allow immediate complementation of P21 function. Complementation is instead restored if specific small deletions in the 3’-UTR region occurs: such deletions likely occur through local cell autonomous replication, since the mere transcription of RNA from the host RNA polymerase guided by 35S promoter would not result in >6 base long deletions (Sydow and Cramer, 2009), while recombination occurring from viral replication would more easily result in the deletions we observed. Therefore, our data shows that the mere local expression of the protein *in trans* does not complement the P21 defective mutants: a full complementation requires P21 expression in the context of viral replication, and in our case, such complementation requires deletions in the 3’-UTR region for the third RNA. Only through those deletions (that we can mimic with an infectious tripartite system of clones) the system can fully recover the ability to complete the full viral infection cycle, including the ability for transmission to new host plants via whitefly.

A relevant precedent to our experimental approach is the bipartitization of TMV using complementing DI expressing different regions of the genome, but in that case no specific adjustment was required (or tested for), and the resulting bipartite tobamovirus was tested in plants only for local infection (Knapp et al., 2005; Lewandowski and Dawson, 2000).

Another relevant example is the case of the first deconstructed virus vector assembled for cowpea mosaic virus (CPMV): in a similar approach to ours, but for the purpose of transforming CPMV in an expression vector, RNA2 was split in two defective RNAs, one coding for the MP (fused to GFP) and one coding for the CPs (L and S proteins): such vector was unstable and recombined to form a full-length RNA2, reverting to bipartite: in our case, the tripartite virus was stable and never reverted to the WT bipartite form (Verver et al., 1998).

To determine whether the deletions we observed had an effect on the secondary structure of its derived RNAs, we compared the predicted secondary RNA structure of the derived RNAs with 6-bp, 9-bp and 61-bp deletions in the R2-p21 3’-UTRs and found out that the absence of these 9bp- and 61-bp would change the conformation of the 3’-UTR of the derived RNAs, but not in the case of the 6-bp deletion. Deletions at the 3’-UTRs of torradoviruses have only been naturally reported for the RNA1 of the Polish tomato torrado virus isolate Kra (Budziszewska et al., 2014); nevertheless, the fact that the 6-bp deletion does not result in major changes in predicted RNA secondary structure seem to rule out the possibility that 3’-UTR conformational changes are necessary for the replication of the RNA3 in the tripartite virus context.

To investigate whether these deletions would have an effect on the biological fitness of the arisen tripartite viruses, we engineered two constructs harboring the 9- and 61-bp deletions and confirmed that the tripartite viruses were able to replicate in *N. benthamiana* plants and in both cases, they were stable in the population for the eleven passages we assayed. In addition, the biological host range was similar for ToANV-WT and ToANV-TripB when agroinfiltrated in different hosts. We did note that in tomato plants systemic symptoms were milder for ToANV-TripB. Possibly, systemic accumulation of the ToANV-TripB in this host was not enough to reproduce efficient mechanical inoculation on tomato which in turn reduces the hazard since ToANV-TripB would not infect efficiently tomato plants in nature.

Our newly originated tripartite virus was selected in *N. benthamiana* plants and, a fitness cost was evident when moving this virus in tomato through, mechanical inoculation: in fact, mechanical inoculation from tomato to tomato plants of TripB did not produce infection in tomato plants suggesting a fitness cost of the *N. benthamiana*-evolved TripB in tomato plants. In addition, when competition experiments of ToANV WT vs TripB in *N. benthamiana* plants were performed, the bipartite WT virus outcompetes the TripB virus, suggesting also in this host a fitness cost.

ToANV TripB was also capable of forming virions and being transmitted by its whitefly vector, and allowed the use of this tripartite virus to be engineered as a viral vector to study in the future the role of different P21 conserved motifs on torradovirus biology. The engineered viral vector was able to infect *N. benthamiana* systemically.

Transitions from monopartite to multipartite viral forms have been described *in vitro* under conditions of high multiplicity of infection (MOI) for animal viruses. Passages of foot-and-mouth disease virus (FMDV) in cell culture generated defective RNAs that were infectious by complementation (García-Arriaza et al., 2004). Their demonstration indeed relied on infectious clones that split the original genome in two segments, mimicking the bipartite virus obtained *in vitro* from natural evolution. In that case, the bipartite system had better fitness (at high MOI) because of increased particle stability of virions with smaller genomic RNA (Ojosnegros et al., 2011) contrary to what happens in our system, that has lower fitness and cannot extinguish the WT version of RNA2 in co-inoculation experiments, showing a basic difference between a natural evolutionary adaptation to multipartitism (*in vitro*) and the one forced by our experimental design through infectious clones.

Here using knowledge derived from the tripartite adaptation studied we provide a ToANV GFP expression vector that can infect systemically plants with a deconstructed-vector approach. Secovirid-based systems for the expression of whole genes have been based on modifications of the RNA2 and co-inoculation with unmodified RNA1 to provide the proteins required for replication and processing (Choi et al., 2019; Zarzyńska-Nowak et al., 2024). Some of the main constraints are related to: i) genome size: by introducing non-viral sequences into the viral genome can result difficult for icosahedral viruses subjected to constraints of genome size for efficient encapsidation; ii) duplication of cleavage sites: the cleavage sites of the encoded polyproteins have to be identified and the duplication of these cleavage sites can lead to recombination or gene products with bigger size. One of the alternatives to modify the RNA2 is creating deconstructed viruses with more than two RNA molecules. In the case of torradoviruses, two viral vectors tagged with GFP had been engineered by inserting the GFP between the MP and the coat protein VP35 and duplicating the cleavage sites but while ToTV was successful our ToANV failed to reproduce systemic infection (Ferriol et al., 2016b; Wieczorek et al., 2020) .

Evolution of multipartitism is an interesting aspect of virus evolution that has received some theoretical and experimental attention (Leeks et al., 2023): furthermore, the more detailed evolutionary history of viruses inferred from recent description of the virosphere shows more and more examples of closely related monopartite viruses that have close relative bipartite viruses (Edgar et al., 2022; Neri et al., 2022). Some examples are the case of the genera closterovirus and crinivirus in the *Closteroviridae* family (Fuchs et al., 2020) or a mix of monopartite and bipartite genomes in the genus *Penicillimonavirus* in the *Mymonaviridae* family (Pagnoni et al., 2023). Often, modelling mathematically the natural transition from monopartitism to multipartitism takes in consideration the fact that shorter RNA are easier to replicate, but that any non-cell autonomous element of the virus life cycle is complicated by mutli-segmentation, for example, requiring all the segments to be coordinately transported to a neighboring cells. Here we show that indeed evolution from a bisegmented to trisegmented virus originates a stable virus, but with lower fitness in the original host: possibly this lower fitness in the original host can turn into new adaptive features when moving to a different host.

Finally, the results obtained in this work are important to understand the role of the P21 protein in the infection cycle of torradoviruses. We foresee the use of the engineered viral vector to study the still unresolved molecular and biochemical properties of P21 in the torradoviruses infectious cycle.

## MATERIALS AND METHODS

### Plasmid constructs

Previously, a series of ToANV constructs derived from ToANV infectious cDNA clones were engineered to study the role of RNA2-ORF1 protein in ToANV infections (Ferriol et al., 2018, 2016b) and were used in this study. We generated two constructs that would not express the RNA2-ORF1 protein (R2-p21-sc and R2-Δp2, (Ferriol et al., 2018)) by using two different strategies: inserting three stop codons or engineering a deletion of 190 aas in the RNA2-ORF1 coding sequence (R2-p21-sc and R2-Δp21, respectively). In addition, we engineered a construct that will only express the RNA2-ORF1 protein and was flanked by the original RNA2 5’- and 3’-UTRs (R3-p21) (Ferriol et al., 2018).

To obtain an RNA2-ORF1 construct with silent mutations in the RNA2-ORF1 sequence, an inverse PCR was performed using R3-p21 and the oligonucleotides ToANV-rectest-For/ToANV-rectest-Rev (Supplementary Table 1) using Phusion High-Fidelity DNA Polymerase (New England BioLabs) following manufacturer’s instructions. The resulting plasmid was named R3-p21-rc.

To generate two constructs with 9-nt and 61-nt deletions in the 3’-UTR of the R3-p21 construct, two inverse PCRs were carried out using R3-p21 as a template and oligonucleotides Trip.A.For/Trip.A.Rev and Trip.B.For/Trip.B.Rev (Supplementary Table 1) using Phusion High-Fidelity DNA Polymerase (New England BioLabs) following manufacturer’s instructions. The resulting plasmids were named R3-p21-TripA (9-bp deletion) and R3-p21-TripB (61-nt deletion).

In order to obtain an RNA3 that expressed GFP we engineered an *Nco*I site in frame with the MP coding sequence, but just downstream of the termination of the RNA2-ORF1 coding sequence with an inverse PCR using Phusion High-Fidelity DNA Polymerase with oligonucleotides RNA2-ORF1-723-F (phosphorylated at its 5’ end) and RNA2-ORF1-NcoI-Rev (Supplementary Table 1) and using as template R3-p21-TripB. The resulting amplification product was digested with *Nco*I. A GFP fragment was amplified with oligonucleotides GFP-NcoI-F and GFP-SmaI-R (Supplementary Table 1) from a GFP encoding plasmid previously described (Crivelli et al., 2011) and digested with *Nco*I and *Sma*I. The two digested PCR product (vector and GFP insert) were ligated to originate the plasmid R3-p21-TripB-GFP.

### Agroinfiltration assays

Plasmids were transformed into *Agrobacterium tumefaciens* strain C58C1 cells and infiltrated into *N. benthamiana* or tomato line cv “York” as previously described (Ferriol et al., 2016b). Equal volumes of *A. tumefaciens* cells harboring the different constructs were mixed prior to infiltration and used at an optical density (OD) at 600nm of 1.

### Detection by DAS-ELISA

DAS-ELISA was carried out as previously described (Ferriol et al., 2018). Briefly, crude extracts from 0.5 cm disc diameter of plant leaf tissue were homogenized in 1 ml of PBS-Tween buffer containing 2% (w/v) PVP. Specific ToANV-anti-virion antibodies were used (Turina et al., 2007).

### RNA purification and northern blot analysis

Total RNAs were extracted from 100 mg of plant tissue using RNeasy Plant Mini Kit (Qiagen). One μg of total RNA was used for northern blot analysis using two ToANV probes (+ and – strand) that will hybridize to the 3’-UTR of ToANV RNA1 and RNA2 and all RNA2-derived constructs used in this work. Northern blot analysis, probe synthesis, hybridization and washes were performed as previously described (Ferriol et al., 2016b).

### RT-PCR and sequencing

Sanger sequencing of recombinant and mutagenized plasmids and of purified RT-PCR amplification products was carried out by Biofab Research (Rome, Italy). RT-PCR was carried out from total RNA extracted as detailed above; cDNA was synthesized with RevertAid First Strand cDNA Synthesis Kit (ThermoScientific) reverse transcription kit and PCR was carried out using OneTaq® DNA Polymerase reagents (NEB) using conditions previously detailed and oligonucleotides described in Suppl. Table 1 and in (Ferriol et al., 2018).

### Serial passages and competition experiments

For both these experiments, *N. benthamiana* plants grown in an insect-proof air-conditioned greenhouse (26°C ± 1°C) under natural light conditions. For the serial passage experiment, agroinfiltration was performed as above to obtain systemically infected plants (three plants for each treatment), with WT and TripB clones (R1-WT+R2-WT and R1-WT+R2-Δp21+R3-p21-TripB, respectively). After the initial inoculation by agro-infiltration a number of subsequent serial mechanical inoculations of plants dusted with carborundum were performed using as source of inoculum a mix of portions of three leaves for each treatment (1 g of fresh weight) from infected plants 10 dpi, homogenized using mortar and pestle in 5 mL of inoculation buffer (50 mM potassium phosphate buffer, pH 7, containing 1 mM Na-EDTA, 5 mM sodium diethyldithiocarbamate and 5 mM sodium thioglycolate) as previously detailed (Bertran et al., 2016). Serial inoculations were performed for eleven times.

For the competition experiment, *N. benthamiana* plants were agroinfiltrated with two mixes of *A. tumefaciens* cells harboring the desired clones in equal amounts (each suspension 1OD): i) R1-WT+ R2-WT+pJL89 (empty vector) and ii) R1-WT+R2-Δp21+R3-p21-TripB. Systemically infected leaves were harvested 5 dpi and processed for northern blot analysis as described above.

### Virion purification, western blot and TEM observations

Virion purification was performed through differential centrifugation steps and sucrose gradients exactly as previously described (Turina et al., 2007). Sodium dodecyl sulfate-polyacrylamide gel electrophoresis (SDS-PAGE) and western blot analysis of purified preparations post sucrose gradients were carried out as described previously in detail (Rastgou et al., 2009) using ToANV polyclonal antibodies raised against purified virions at 1:2000 dilution (Turina et al., 2007). Transmission electron microscopy (TEM) observation of purified particles through negative staining with uranyl acetate was done as previously detailed (Ferriol et al., 2018). Electron dense and electron transparent particles were counted in different areas of TEM grids, reaching a total of about 600 viral particles.

### RNA secondary structure analyses

The secondary structures of the 3’-UTRs of the RNA2 WT, R2-p21 and R2-p21-6bp-, R2-p21-9bp (TripA) and R2-p21-9bp (TripB) derived RNAs were predicted using the Mfold program at the website (http://unafold.rna.albany.edu/?q=mfold/RNA-Folding-Form) (Zuker, 2003), and illustrated with the RNA2Drawer program (Johnson et al., 2019).

### Total RNA extraction and RNA-seq transcriptome assembly

Total RNAs were extracted using Total Spectrum RNA reagents (Sigma-Aldrich). RNAs were quantified using a NanoDrop 2000 Spectrophotometer (Thermo Scientific). One ug of RNA pooled from equal amounts of three infected plants from each of the two treatments (WT and TRIP-B) were sent to sequencing facilities (Macrogen, Seoul, Rep. of Korea). Ribosomal RNA (rRNA) was depleted (Ribo-ZeroTM Gold Kit, Epicentre), cDNA libraries were built with TrueSeq total RNA sample kit (Illumina) and sequencing was performed by an Illumina HiSeq4000 system generating paired-end sequences. Bioinformatic analyses of RNAseq raw reads was performed as previously described. BWA (0.5.9) (Li and Durbin, 2009) and SAMtools (0.1.19) (Li et al., 2009) were used to map reads on reference sequences of the infectious clones and visualized with IGV (Thorvaldsdóttir et al., 2013). Only mutations present in more than 2% of the reads were scored.

### Whitefly transmission experiments

All plants were grown in 10-cm diameter pots in a growth chamber at 26°C under a 14-h light regime or in insect-proof greenhouses under natural light conditions. *Trialeurodes vaporariorum* whiteflies from a population harvested from a greenhouse at the Turin botanical garden were maintained on melon plants in insect proof growth chambers at 26°C ± 1°C, with a 16:8 h (1ight- dark) photoperiod and passaged to new healthy plants every two weeks. Non viruliferous *Trialeurodes vaporariorum* whiteflies were then transferred to ToANV-WT or ToANV-TripB-infected *N. benthamiana* plants by gentle shaking as previously described and removed after an acquisition access period (AAP) of 48 hr (Ferriol et al., 2016b). For inoculation access, adult whiteflies were placed on receptor plants (*Physalis floridiana* and *Nicotiana occidentalis*) in glass tubes (6 cm diameter, 12 cm high), topped with a nylon screen and after an inoculation access period of 72h plants were sprayed with insecticide to remove whiteflies; after that plants were transferred back to the greenhouse and monitored for symptoms and for sample harvesting for two weeks.

### GFP visualization

*N. benthamiana* leaves were agroinfiltrated with a combination of *A. tumefaciens* cells harbouring R1-WT+R2-Δp21+R2-p21-TripB-GFP and then examined for GFP fluorescence at 3- and 6-days post agroinfiltration. Plants agroinfiltrated with R1-WT+R2-Δp21+R2-p21-TripB were used as a negative control. Fluorescence was observed using a Leica M205 FA stereo microscope (Leica Microsystems, Wetzlar, Germany). A 488-nm Argon laser line and a window of 500 to 525 nm were used for excitation and for collection of the GFP signal, respectively. Pictures were captured using a ×10 or ×20 objective and processed (for trimming, contrast and brightness adjustment, and figure assembly) using the GNU Image Manipulation Program (https://www.gimp.org/).

## Data availability

Raw reads from the Illumina sequencing are available upon request

## Author Contribution

Conceptualization: MT, IF Funding Acquisition: MT, IF, BF writing review and editing: MT, IF, BF, MC, MV, LN Writing original draft: MT and IF, Data curation LN, Investigation Methodology, MC, MV, MT, IF

## Acknowledgements

This research was partially funded by grant SCB11058 from the California Department of Food and Agriculture. M.T. was supported by a short-term mobility scholarship from the Italian CNR . I.F. was supported by a research grant from the Federation of European Microbiological Society (FEMS-RG-2016-0120). We also would like to acknowledge the technical support of Riccardo Lenzi.

## ONLINE SUPPLEMENTARY MATERIAL

**Supplementary Table 1.** Oligonucleotides used in this study.

**Supplementary Table 2.** Nucleotide and amino acid changes in RNA1 of tomato apex necrosis virus WT and Trip B after eleven passages in *Nicotiana benthamiana* and tomato plants.

**Supplementary Table 3.** Nucleotide and amino acid changes in the RNA2 of tomato apex necrosis virus (ToANV) WT and Trip B lineages after eleven passages in *Nicotiana benthamiana* and tomato plants.

**Supplementary Table 4.** Nucleotide and amino acid changes in the RNA3 of tomato apex necrosis virus (ToANV) and Trip B lineages after eleven passages in *Nicotiana benthamiana* and tomato plants.

**Supplementary Figure 1. Analyses of the progeny of upper non-inoculated leaves of plants agroinfiltrated with *A. tumefaciens* harboring the R3-p21-rc shows evidence of recombination between R3-p21-rc and R2-p21sc.** Sequence alignments of cDNA obtained through RT-PCR across spanning the initial coding region of p21 present in RNA2 of Tomato apex necrosis virus. Systemic infection resulting from the attempt at complementing *in trans* the knockout of the p21 protein obtained through three stop codons downstream of the p21 AUG in the RNA2 was characterized by sequencing the RT-PCR products from RNA extractions. Here using a p21 construct that carries a marker mutation (R3-p21-rc) we show that the progeny of virus that systemically infects the plants 2 weeks post agro-inoculation carries the marker in the RNA2 (RecTest2, RecTest3 and RecTest4); at the same time the third RNA provided in trans is not present. Underlined is the p21 coding sequence in the R2-WT construct. Dots represent identical nucleotides in the alignment.

**Supplementary Figure 2. Comparison of the predicted secondary structures of the 3’-UTRs of ToANV RNA2-WT, R3-p21 with the R3-p21 derived RNAs with 6-, 9- and 61-bp deletions.** Nucleotides that have been deleted are indicated in blue. Nucleotides placed before the deletion are indicated in black and nucleotides placed after the deletion are indicated in yellow.

**Supplementary Figure 3. Multiple nucleotide alignment of the RNA1 and RNA2 3’-UTRs of different ToMarV isolates in comparison with the R3-p21 with the R3-p21 derived RNAs with 6-, 9- and 61-bp deletions.** The 6-, 9- and 61-bp deletions are indicated in yellow. GenBank accession numbers for virus isolates are: ToMarV isolate Ahome (RNA1: MK733734 and RNA2: MK726319), ToMarV isolate PRI-TMarV0601_(RNA1: EF681764 and RNA2: EF681765), ToANV isolate VE434 (RNA1: EF063641 and RNA2: EF063642), ToNDV isolate R (RNA1: KC999058 and RNA2: KC999059).

**Supplementary Figure 4. Systemic symptoms of tomato plants (cv YORK) inoculated with WT tomato apex necrosis (R1-WT+R2-WT) and with the tripartite B infectious clone combination (R1-WT+R2-Δp21+R3-p21-TripB).** Pictures were taken two weeks post agroinfiltration. Strong deformation of the systemically infected leaf can be seen with the WT clone combinations, while milder symptoms are displayed with the tripartite infectious clone combination.

**Supplementary Figure 5. Northern blot analysis of RNA extracts from un-inoculated leaves of *Nicotiana occidentalis* inoculated with insects that acquired the virus on systemically infected *N. benthamiana* plants with the WT (WT-Insect) and TripB tripartite virus (lane TripB-Insect A and TripB-insect B).** Lane labelled TripB is the original TripB infected *N. benthamiana* plant used for acquisition. Membranes were exposed to films for 12 hrs (left panels) and 48 hrs (right panels). Arrows point to the three genomic RNAs. Lower panels represent rRNA loadings of the same membranes pre-stained with methylene blue. A negative sense riboprobe was used hybridizing with positive sense viral RNA.

