## Supplementary file for "Monitoring the directed evolution to a tripartite genome from a bipartite torradovirus genome"

**Supplementary Table 1.** Oligonucleotides used in this study**.**

| Primer name | Sequence (5'-3') | Objective |
|---|---|---|
| **ToANV-Rectest-For** | GG**a**CACTTAAATACATCAGTAGAGGAGC | Engineer R2-p21-rc |
| **ToANV-Rectest-Rev** | TATAAATGACATGAAAATGTTACTCAAA | |
| **Trip-A-For** | TCCCCTCTTCTTCCCTAGGT | Engineer R2-p21-TripA |
| **Trip-A-Rev** | TCATCACTACCCCTAATAGTGAC | |
| **Trip-B-For** | TTCCCTAGGTTCGTCCAGAGG | Engineer R2-p21-TripB |
| **Trip-B-Rev** | GAAATTATCGAATTCCAAACGGC | |
| **ToANV-RNA2-1For-Long** | TTAAAAGAATTTTAATATATATCTGTCA | Sequencing |
| **RNA2-ORF1-723-F** | 5' [phos]-AAAAGTACCATGCTACTGGGTATATAGTAG | Reverse amplification for GFP insertion |
| **RNA2-ORF1-NcoI-Rev** | TCA*CCATGG*TTTTACTGGGGCCCATTTTCAGTCC | Reverse amplification for GFP insertion |
| **GFP-NcoI-F** | AAA*CCATGG*TGAGCAAGGGCGAGGAG | GFP amplification |
| **GFP-Sma-R** | AAA*CCCGGG*TTACTTGTACAGCTCGTCCATG | GFP amplification |
| **5'-UTR-For** | ACGTCGTTGTGCGATCTACG | Sequencing Tripartite |
| **R2-ORF2-Rev** | GCTTGTGCACCATCAGCC | Sequencing Tripartite |

The mutation in the construct R2-p21-rc is indicated in red and bold. Restriction sites used for cloning are indicated in italics and grey.

**Supplementary Table 2**. Nucleotide and amino acid changes in RNA1 of tomato apex necrosis virus (ToANV) WT and Trip B after eleven passages in *Nicotiana benthamiana* and tomato plants.

| Sample | RNA | HOST | # reads | Average coverage | Genomic Region (Nucleotide changes) | | |
|---|---|---|---|---|---|---|---|
| | | | | | **5'-UTR** | **R1-ORF1** | **3'-UTR** |
| **WT Spring 2016** | R1 | *N. benthamiana* | 141,346 | 3,654 | No changes | No changes | C6796T (79%C; 21%T)<br>G6839A (40%G; 55%A)<br>C6859T (27%C; 73%T)<br>T6876C (28%T; 72%C)<br>A6896G (30%A; 70%G)<br>C6943T (46%C; 54%T)<br>G6953A (73%G; A27%) |
| **WT September 2016** | R1 | *N. benthamiana* | 2,116,465 | 29,481 | No changes | **G2257A (29%G; 71%A); R705K** | G6839A (54%G; 46%A)<br>C6859T (37%C; 63%T)<br>T6876C (38%T; 62%C)<br>A6896G (40%A; 60%G)<br>C6943T (56%C; 44%T) |
| **Tripartite B September 2016** | R1 | *N. benthamiana* | 460,988 | 6,421 | No changes | No changes | G6839A (46%G; 54%A)<br>C6859T (35%C; 65%T)<br>T6876C (35%T; 65%C)<br>C6943T (61%C; 39%T) |
| **Tripartite B September 2016 Tomato** | R1 | tomato | 8,138 | 113 | No changes | No changes | G6839A (48%G; 52%A)<br>C6859T (31%C; 68%T)<br>T6876C (35%T; 65%C)<br>A6896G (36%A; 63%G)<br>C6943T (55%C; 45%T) |

Non-synonymous changes are in bold and red ink. R1: KT756876; R2: KT756877

**Supplementary Table 3**. Nucleotide and amino acid changes in the RNA2 of tomato apex necrosis virus (ToANV) WT and Trip B lineages after eleven passages in *Nicotiana benthamiana* and tomato plants.

| **Sample** | **RNA** | **HOST** | **# reads** | **Av. cov.** | **Genomic Region (Nucleotide changes)** | | | |
|---|---|---|---|---|---|---|---|---|
| | | | | | **5'-UTR** | **R2-ORF1** | **R2-ORF2** | **3'-UTR** |
| **WT Spring 2016** | R2 | *N. benthamiana* | 311,622 | 6,394 | No changes | No changes | No changes | No changes |
| **WT September 2016** | R2 | *N. benthamiana* | 3,623,160 | 74,348 | No changes | **T328A (3%T; 97%A) S60T** | T2182C (0%T; 100%C) | T4532C (78%T; 22%C)<br>C4549T (78%C; 22%T)<br>G4569A (80%G; 20%A) |
| **Tripartite B September 2016** | R2-∆p21 | *N. benthamiana* | 571,330 | 13,144 | No changes | n/a | **G3719A (73%G; 27%A) TGA to TAA (Stop>Stop)** | C4017T (80%C; 20%T) |
| **Tripartite B September 2016** | R2-∆p21 | tomato | 14,330 | 329 | No changes | n/a | **G3719A (75%G; 25%A) TGA to TAA (Stop>Stop)** | No changes |

Non-synonymous changes are in bold and red ink. N/a: not available because this region is not present.

**Supplementary Table 4**. Nucleotide and amino acid changes in the RNA3 of tomato apex necrosis virus (ToANV) and TripB lineages after eleven passages in *Nicotiana benthamiana* and tomato plants.

| **Sample** | **RNA** | **HOST** | **# reads** | **Av. cov.** | **Genomic Region (Nucleotide changes)** | | | |
|---|---|---|---|---|---|---|---|---|
| | | | | | **5'-UTR** | **R2-ORF1** | **R2-ORF2** | **3'-UTR** |
| Tripartite B September 2016 | R3-Trip-B | *N. benthamiana* | 150,721 | 11,433 | No changes | No changes | n/a | Deletion of T1063; 90% of reads have this deletion. 10% of reads without deletion, but T1063C (65%T; 35%C, this is from the 10% without deletion) |
| Tripartite B September 2016 | R3-Trip-B | tomato | 2,809 | 212 | No changes | A722G (76%A; 24%G); TAA to TGA (Stop to Stop) | n/a | No changes |

N/a: not available because this region is not present.

### Supplementary Fig. 1

```
#R2-WT-KT756877   CCAACTTTTGAGTAACATTTTCATGTCATTTATAGGTCACTTAAATACATCAGTAGAGGAGCAAGCATTT
#R3-p21-rc        ....................................A.................................
#RecTest4_1For    ....................................A.................................
#RecTest3_1For    ....................................A.................................
#Rec-Test2_1For   ....................................A.................................
```

**Supplementary Figure 1. Analyses of the progeny of upper non-inoculated leaves of plants agroinfiltrated with *A. tumefaciens* harboring the R3-p21-rc shows evidence of recombination between R3-p21-rc and R2-p21sc.** Sequence alignments of cDNA obtained through RT-PCR across spanning the initial coding region of p21 present in RNA2 of Tomato apex necrosis virus. Systemic infection resulting from the attempt at complementing *in trans* the knockout of the p21 protein obtained through three stop codons downstream of the p21 AUG in the RNA2 was characterized by sequencing the RT-PCR products from RNA extractions. Here using a p21 construct that carries a marker mutation (R3-p21-rc) we show that the progeny of virus that systemically infects the plants 2 weeks post agro-inoculation carries the marker in the RNA2 (RecTest2, RecTest3 and RecTest4); at the same time the third RNA provided in trans is not present. Underlined is the p21 coding sequence in the R2-WT construct. Dots represent identical nucleotides in the alignment.

Fig. 2

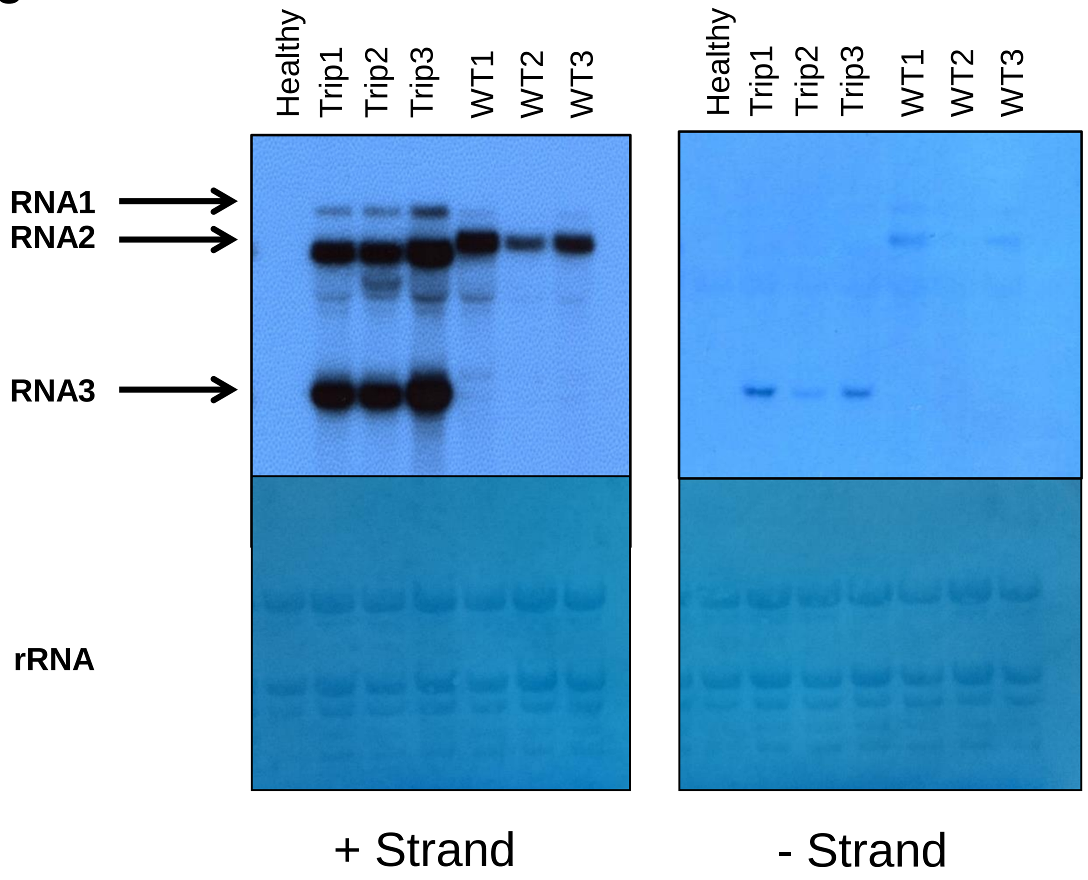


**Supplementary Figure 2. Comparison of the predicted secondary structures of the 3'-UTRs of ToANV RNA2-WT, R3-p21 with the R3-p21 derived RNAs with 6-, 9- and 61-bp deletions.** Nucleotides that have been deleted are indicated in blue. Nucleotides placed before the deletion are indicated in black and nucleotides placed after the deletion are indicated in yellow.

### Supplementary Fig. 3

```
R2-p21                     A--AAGTACCATGCTACTGGGTATATAGTAGCCAAATACACTTATCTACGTGTACTGTTTGTAGTAGCT-
R3-p21-TripA               .--..................................................................-
R3-p21-TripB               .--..................................................................-
R3-p21-6bp_deletion        .--..................................................................-
R2-ToMarV_Ahome            .--.........................................C..............A.......T.-
R2-ToMarV_PRI-TMarV0601    .A-........................................................A.........-
R2-ToANV_VE434             .--..................................................................-
R2-ToNDV_R                 .--...........C........A.....G..........T...A......................-.T
R1-ToMarV_Ahome            GTC.G.C.TT...--...CAA.-.-.-...-.--G...TG.--..----..........G...A.-.-.-
R1-ToMarV_PRI-TMarV0601    GTC.G.C.TT...--...CAA.-.-.-..-..--G...TG.--..----..........A...A.-.-.-
R1-ToANV_VE434             GTC.G.C.TT...--...CAA.-.-.-..G-.--G...TG.--..----..........A...A.-.-.-
R1-ToNDV_R                 GTC.G.C.TT...--...CA..---.T..G--..---...T--G--..T..........---....A-.-

R2-p21                     A-AGTGGTGTTTGCAT--A-T-A-T--G-T-T---TTATTTAAGGGAGAGTACTATCTTTTATTAGATGG
R3-p21-TripA               .-..............--.-.-.-.--.-.-.---...................................
R3-p21-TripB               .-..............--.-.-.-.--.-.-.---...................................
R3-p21-6bp_deletion        .-..............--.-.-.-.--.-.-.---...................................
R2-ToMarV_Ahome            .-...........TT.--G-C-.-.----.-.-A-C.-................G......G........
R2-ToMarV_PRI-TMarV0601    .-...........TT.GC.A.G.-.AA.GG-A---GAG..-.............G.......A.......
R2-ToANV_VE434             .-............----.-.-.-.AT.-.-.---...................................
R2-ToNDV_R                 .GG-..C......TT---.T.-.C.A-.G.A.TAA-.G..-.............G.....CG.....C..
R1-ToMarV_Ahome            .T.-.--C...A.-----G---.-.G-.-.-.---..G..-.............G......G........
R1-ToMarV_PRI-TMarV0601    .------.A.C-.TT---.G-G.-.--.G.-.---..G..-.............G.......A.......
R1-ToANV_VE434             .T.-.--C.C.A.-----G---.-.G-.-.-.---..G..-.............G......G........
R1-ToNDV_R                 .C.-..T....-.T----.---.-.--.A.AC-AAC.T..-............C......C.........

R2-p21                     GAGTCCCCTCCACTATATGGAGACCGATGGATCTACATCGGTATAGAGGTTTGCACGCTCCTCTTT-AAA
R3-p21-TripA               ..................................................................-...
R3-p21-TripB               ..................................................................-...
R3-p21-6bp_deletion        ..................................................................-...
R2-ToMarV_Ahome            ..............CC.........AG..A........T...........................-...
R2-ToMarV_PRI-TMarV0601    ..............TC.........A...A........T...........................-...
R2-ToANV_VE434             ..............T..........A...A.A......T.....G.T.A..A..GT...T.A.C..-...
R2-ToNDV_R                 ............T.C..........AG..A.......CT....C....A.........CT......T...
R1-ToMarV_Ahome            ..............T..........A...A.A......T.....G.T.A..A..GT...T.A.C..-...
R1-ToMarV_PRI-TMarV0601    ..............T..........A...A........T...........................-...
R1-ToANV_VE434             ..............T..........A...A.A......T.....G.T.A..A..GT...T.A.C..-...
R1-ToNDV_R                 ............T.C...........G..A.......CT.........A.........CT.....C-...

R2-p21                     CAAG-ATTCGTGTGCCCTGCTTGGTTAGAAAGTATGTGGTGATTAACACTACTCATTTGGAGTATAAACA
R3-p21-TripA               ....-.................................................................
R3-p21-TripB               ....-.................................................................
R3-p21-6bp_deletion        ....-.................................................................
R2-ToMarV_Ahome            ....T..-..............................................................
R2-ToMarV_PRI-TMarV0601    T...-T..........................C.....................G............G..
R2-ToANV_VE434             T.T.-G...AC.............C..........................................G..
R2-ToNDV_R                 T.T.-G.......A..........C.......C.C......G............................
R1-ToMarV_Ahome            T.T.-G...AC.............C.............................G............G..
R1-ToMarV_PRI-TMarV0601    T...-T..........................C.....................G............G..
R1-ToANV_VE434             T.T.-G...AC.............C..........................................G..
R1-ToNDV_R                 ..T.-G..........................C.C......G.C..........G...............

R2-p21                     ATAGACCTCATGATGTCTAACTCATGCGTGATTGCTCATGTACGAAATAAATGAGCCGTTTGGAATTCGA
R3-p21-TripA               ......................................................................
R3-p21-TripB               ......................................................................
R3-p21-6bp_deletion        ......................................................................
R2-ToMarV_Ahome            ......................................................................
R2-ToMarV_PRI-TMarV0601    .................................................................C....
R2-ToANV_VE434             .................C................T...................A...............
R2-ToNDV_R                 ......T..............................G........G....C...T..............
R1-ToMarV_Ahome            .................C................T...................A...............
R1-ToMarV_PRI-TMarV0601    .................................................................C....
R1-ToANV_VE434             .................C................T...................A...............
R1-ToNDV_R                 ..........................T..........G....T...G....C...T..............

R2-p21                     TAATTTCCCTTAGCTGCTGAACAGCTGTCACTATTAGGGGTAGTGATGAAGTTGTAGATCCCCTCTTCTT
R3-p21-TripA               ................................................---------.............
R3-p21-TripB               .......-------------------------------------------------------------..
R3-p21-6bp_deletion        ................................................------................
R2-ToMarV_Ahome            ...............................C.........G.............G....A.........
R2-ToMarV_PRI-TMarV0601    ........T..........T.........................GC........GAG..........C.
R2-ToANV_VE434             ...............................C.........G..........A..GAGAAA.........
R2-ToNDV_R                 C......................................................GTCAAA...T..T..
R1-ToMarV_Ahome            ...............................C.........G..........A..GAGAAA.........
R1-ToMarV_PRI-TMarV0601    ...................C.........................GC.....C..GAG..........C.
R1-ToANV_VE434             ...............................C.........G..........A..GAGAAA.........
R1-ToNDV_R                 .......................................................GTCAAA.........
```

```
R2-p21                    CCCTAGGTTCGTCCAGAGGGTTCAAAGATC--T--ACTTTCTTTGCCAAGAAACAATGAATGGCACGTGT
R3-p21-TripA              ..............................--.--...................................
R3-p21-TripB              ..............................--.--...................................
R3-p21-6bp_deletion       ..............................--.--...................................
R2-ToMarV_Ahome           ..............................--C--.TC.......T......G.........A.......
R2-ToMarV_PRI-TMarV0601   .............................T--C--..C.......T......G.G.......A.......
R2-ToANV_VE434            ...............A.......--....T--CTC..C..............GT..A.............
R2-ToNDV_R                .......................--....TGAC--C.C..T...........GT................
R1-ToMarV_Ahome           ...C...................--....T--CTC..C..............GT..A.............
R1-ToMarV_PRI-TMarV0601   .............................T--C--..C.......T......G.........A.......
R1-ToANV_VE434            .......................--....T--CTC..C..............GT..A.............
R1-ToNDV_R                .......................--....TGAC--..C..............G.................

R2-p21                    TGCGTCGACAAAGCACCGATTC-A-CATGGTTAG-TGGATT-T---TAGTATTATATTAGATTTGTTAGT
R3-p21-TripA              ......................-.-.........-......-.---........................
R3-p21-TripB              ......................-.-.........-......-.---........................
R3-p21-6bp_deletion       ......................-.-.........-......-.---........................
R2-ToMarV_Ahome           ...........T......-...T.-.........-.A....-.---..-...................A.
R2-ToMarV_PRI-TMarV0601   ..................-...T.-.G.......-.A..AAG.---..-.....................
R2-ToANV_VE434            ......A...........--..T.G.GA......C.A..A.T.---..-.............A..AA.A.
R2-ToNDV_R                ..................--..C.G.T.......C....A.T.GTT..-.................C.A.
R1-ToMarV_Ahome           ..................--..T.G.GA......C.A..A.T.---..-.............A..AA.A.
R1-ToMarV_PRI-TMarV0601   ..................-...T.-.G.......-.A..AAG.---..-.....................
R1-ToANV_VE434            ..................--..T.G.GA......C.A..A.T.---..-.............A..AA.A.
R1-ToNDV_R                ...A..............--..C.G.T.......C...GA.T.GTC..-.................C...

R2-p21                    CAATTGTGTGATTTCTT-TCT-TA-ATTTAGAAGGTTTTTCGTGGCGATAAGAAGGGTTTGTCCTTTTAC
R3-p21-TripA              .................-...-..-.............................................
R3-p21-TripB              .................-...-..-.............................................
R3-p21-6bp_deletion       .................-...-..-.............................................
R2-ToMarV_Ahome           .................-...-..G...-..G.A..............C....G................
R2-ToMarV_PRI-TMarV0601   .................-..C-..G...-..G.A.....C..........G...................
R2-ToANV_VE434            TT...............-...G..-G.......A................GAG.................
R2-ToNDV_R                .TG...........T..T.T.-..-G.....GGA...........G....GA...............C..
R1-ToMarV_Ahome           TT...............-...G..-G.......A................GAG.................
R1-ToMarV_PRI-TMarV0601   .................-...-..G...-..G.A.....C..........G...................
R1-ToANV_VE434            TT...............-...G..-G.......A................GAG.................
R1-ToNDV_R                .T...............C...-..-G.....GGA.....C..........GA..................

R2-p21                    CTTCTTTGCTATGCTGGACACAAAAAGATTTTCTTTTCTTTTATTTT
R3-p21-TripA              ...............................................
R3-p21-TripB              ...............................................
R3-p21-6bp_deletion       ...............................................
R2-ToMarV_Ahome           ...T............................T....T.........
R2-ToMarV_PRI-TMarV0601   ...............................................
R2-ToANV_VE434            ..CTC.........................................C
R2-ToNDV_R                ...TC.....G.....................T....T.........
R1-ToMarV_Ahome           ..CTC..........................................
R1-ToMarV_PRI-TMarV0601   ...............................................
R1-ToANV_VE434            ..CTC.........................................C
R1-ToNDV_R                ...TC..........................................
```

**Supplementary Figure 3. Multiple nucleotide alignment of the RNA1 and RNA2 3'-UTRs of different ToMarV isolates in comparison with the R3-p21 with the R3-p21 derived RNAs with 6-, 9- and 61-bp deletions.** The 6-, 9- and 61-bp deletions are indicated in yellow. GenBank accession numbers for virus isolates are: ToMarV isolate Ahome (RNA1: MK733734 and RNA2: MK726319), ToMarV isolate PRI-TMarV0601_(RNA1: EF681764 and RNA2: EF681765), ToANV isolate VE434 (RNA1: EF063641 and RNA2: EF063642), ToNDV isolate R (RNA1: KC999058 and RNA2: KC999059).

Supplementary on line Fig. 4

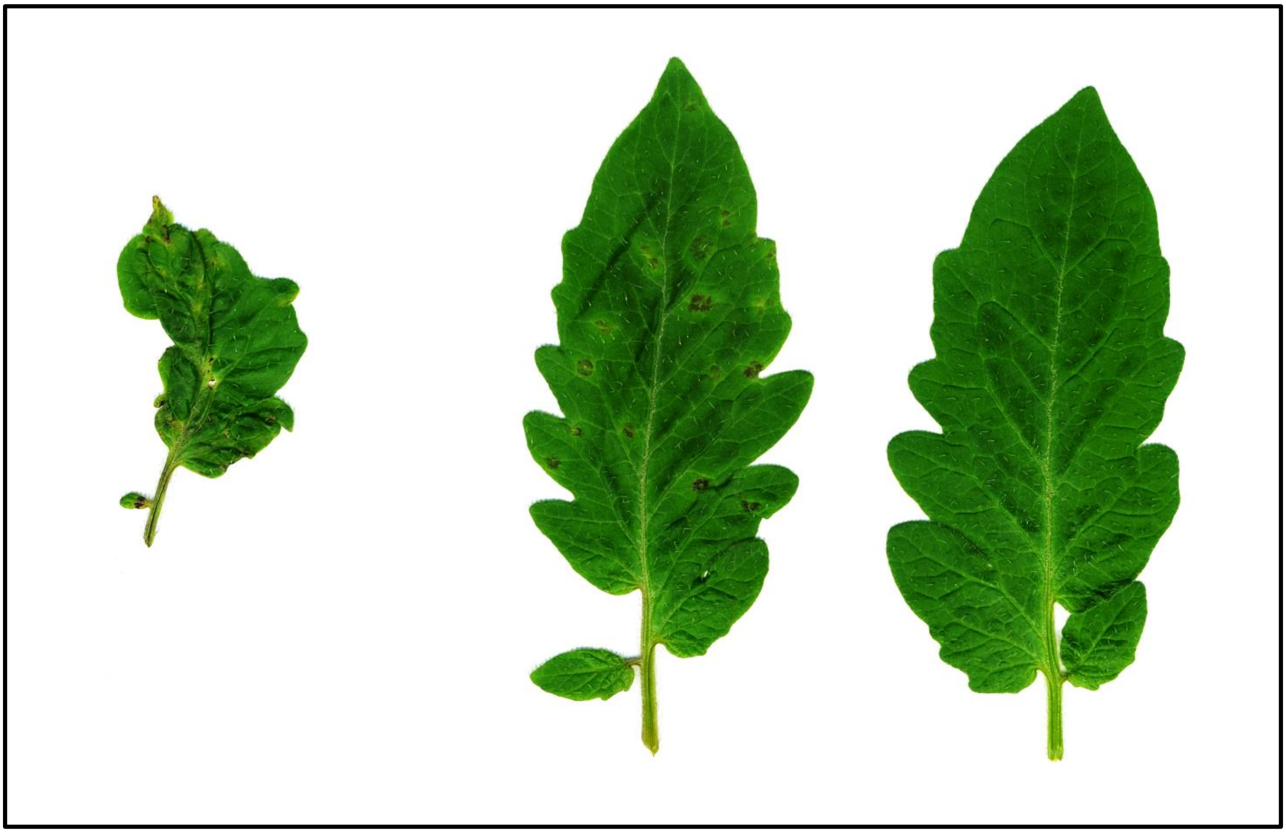

R1-WT+R2- WT

R1-WT+R2-Δp21+R3-p21-TripB

Mock

**Supplementary on line Figure 4:** Systemic symptoms of tomato plants (cv YORK) inoculated with WT tomato apex necrosis (R1-WT+R2-WT) and with the tripartite B infectious clone combination (R1-WT+R2-Dp21+R3-p21-TripB). Pictures were taken two weeks post agroinfiltration. Strong deformation of the systemically infected leaf can be seen with the WT clone combinations, while milder symptoms are displayed with the tripartite infectious clone combination.

### Suppl. Fig. 5

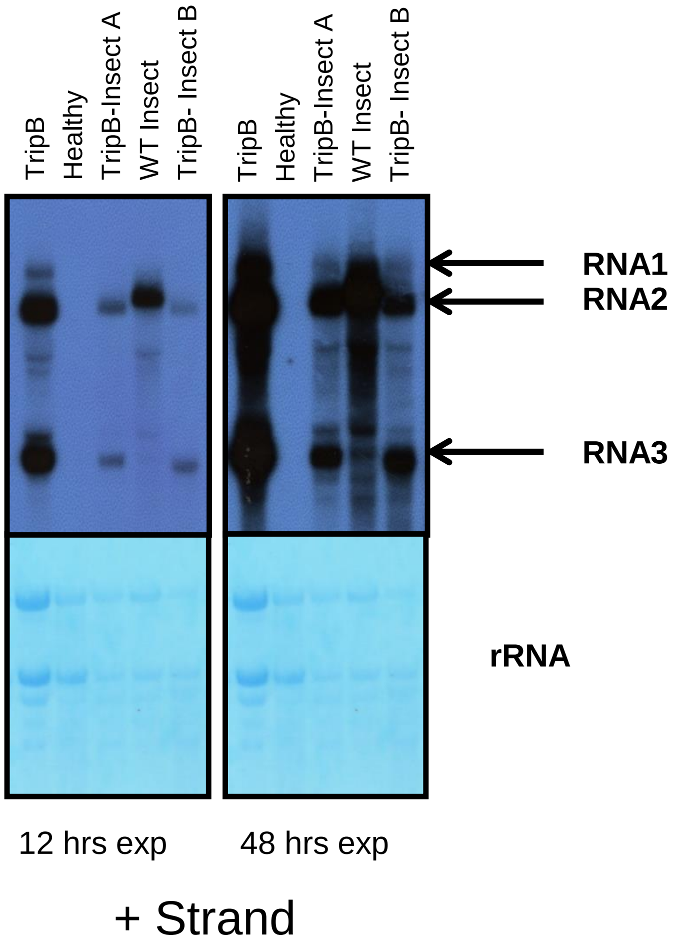


**Supplementary Figure 5. Northern blot analysis of RNA extracts from un-inoculated leaves of *Nicotiana occidentalis* inoculated with insects that acquired the virus on systemically infected *N. benthamiana* plants with the WT (WT-Insect) and TripB tripartite virus (lane TripB-Insect A and TripB-insect B).** Lane labelled TripB is the original TripB infected *N. benthamiana* plant used for acquisition. Membranes were exposed to films for 12 hrs (left panels) and 48 hrs (right panels). Arrows point to the three genomic RNAs. Lower panels represent rRNA loadings of the same membranes pre-stained with methylene blue. A negative sense riboprobe was used hybridizing with positive sense viral RNA.
